# Structures of the multi-domain oxygen sensor DosP: remote control of a c-di-GMP phosphodiesterase by a regulatory PAS domain

**DOI:** 10.1101/2024.07.24.604967

**Authors:** Wenbi Wu, Pankaj Kumar, Chad A. Brautigam, Shih-Chia Tso, Hamid R. Baniasadi, Daniel L. Kober, Marie-Alda Gilles-Gonzalez

**Affiliations:** Department of Biochemistry, University of Texas Southwestern Medical Center, Dallas, TX 75390, USA; Department of Biophysics, University of Texas Southwestern Medical Center, Dallas, TX 75390, USA; Department of Microbiology, University of Texas Southwestern Medical Center, Dallas, TX 75390, USA

## Abstract

The heme-based direct oxygen sensor DosP degrades c-di-GMP, a second messenger nearly unique to bacteria. In stationary phase *Escherichia coli*, DosP is the most abundant c-di-GMP phosphodiesterase. Ligation of O_2_ to a heme-binding PAS domain (hPAS) of the protein enhances the phosphodiesterase through an allosteric mechanism that has remained elusive. We determined six structures of full-length DosP in its aerobic or anaerobic conformations, with or without c-di-GMP. DosP is an elongated dimer with the regulatory heme and phosphodiesterase separated by nearly 180 Å. In the absence of substrate, regardless of the heme status, DosP presents an equilibrium of two distinct conformations. Binding of substrate induces DosP to adopt a single, ON-state or OFF-state conformation depending on its heme status. Structural and biochemical studies of this multi-domain sensor and its mutants provide insights into signal regulation of second-messenger levels.

## Introduction

The DosP protein of *Escherichia coli* is a heme-based sensor of molecular oxygen, O_2_, with a liganded (oxy) ON-state and an unliganded (deoxy) OFF-state ^1–4^. DosP was originally identified as an O_2_ sensor from the resemblance of its heme-binding Per-Arnt-Sim (hPAS) domain to the oxygen sensing region of the FixL proteins that govern rhizobial nitrogen fixation^5^. In FixL, O_2_ ligation to the hPAS domain is coupled to the status of a histidine protein kinase in the same protein^6^. By contrast, DosP couples a ligation event at the hPAS domain to a phosphodiesterase, or EAL domain, that cleaves c-di-GMP to pGpG^1^. Although DosP contains slightly more than 800 residues, together the hPAS at its N-terminus and EAL at its C-terminus account for only half of it: between those two domains in the linear sequence, one finds a PAS-PAC domain and a silent diguanylate cyclase, or EGTQF domain (**Fig. 1a**). Thus, DosP resides at the intersection of the PAS domain-containing proteins, heme-based sensors, and the EAL-type c-di-GMP phosphodiesterases, all of which belong to highly versatile families of signal-transducing multi-domain proteins ^7–9^.

**Figure 1.**
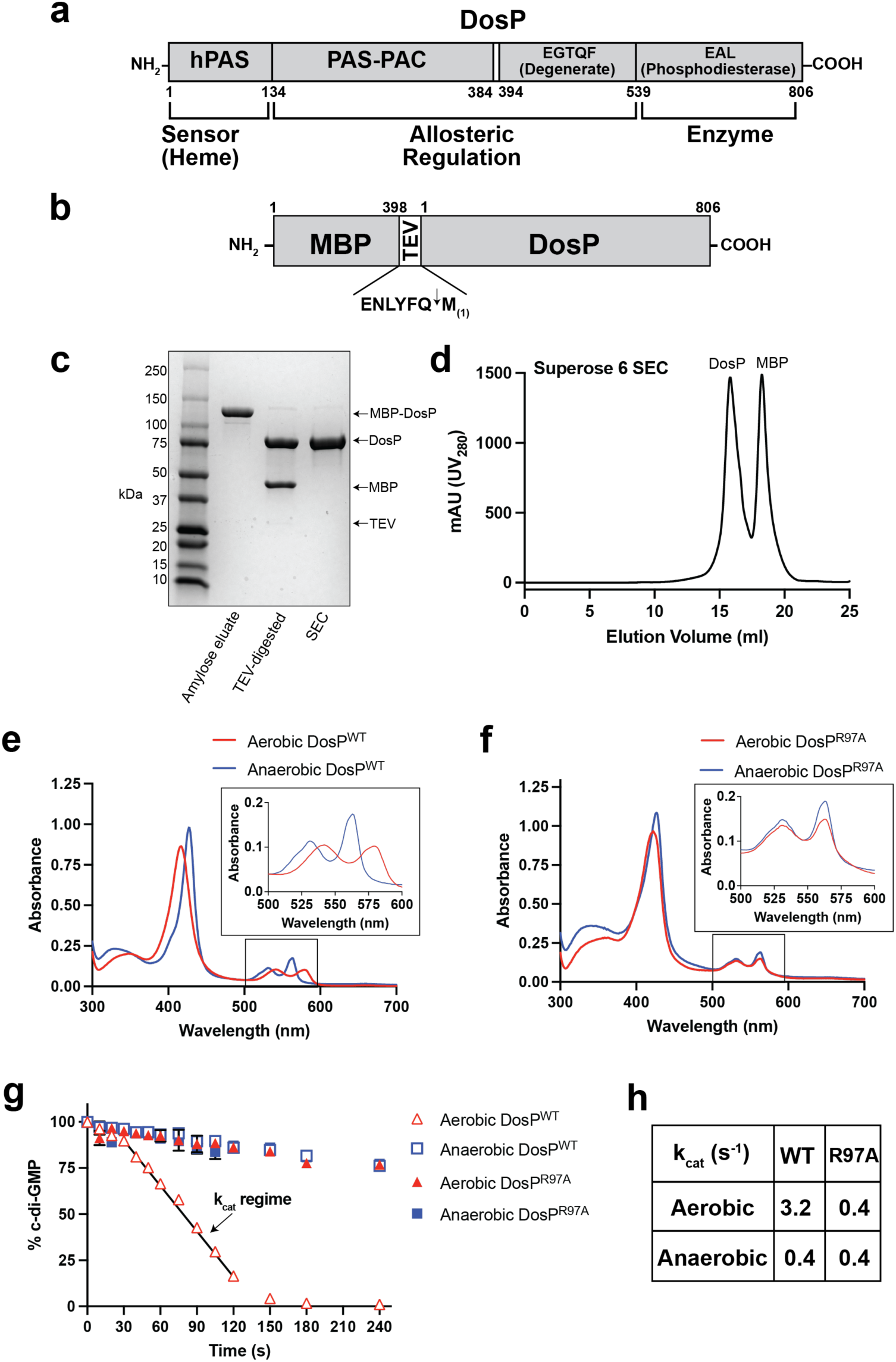
Purification and biochemical characterization of DosP. **a,** Domain organization of DosP over its linear amino acid sequence. **b,** Schematic for the MBP-DosP expression construct. TEV protease recognition sequence is shown under brackets. Cleavage site marked by arrow and resulting N-terminal methionine corresponds to DosP M1. **c,** Coomassie-stained 4-15% SDS-PAGE analysis of purified DosP from the indicated steps. Migration positions of the proteins are indicated with arrows. Molecular weight standards are labeled. **d,** SEC purification of free DosP, MBP, and TEV protease following proteolysis of MBP-DosP using a 10/300 Superose 6 gel filtration column as described in *Methods*. **e,** Absorption spectra of DosP^WT^ under aerobic conditions (red), compared to anaerobic conditions (blue). **f,** Absorption spectra of DosP^R97A^ under aerobic conditions (red), compared to anaerobic DosP^R97A^ (blue). **g,** Turnover of 2 µM c-di-GMP at 25°C by 10 nM protein: aerobic DosP^WT^ (open red triangles), anaerobic DosP^WT^ (open blue squares), aerobic DosP^R97A^ (filled red triangles), and anaerobic DosP^R97A^ (filled blue squares). Black line indicates time regime used to determine k_cat_. the assay buffer was 20 mM Tris-HCl, 2.5 mM MgCl_2_, 2.0 mM DTT, pH 8.0. **h,** k_cat_ values determined from (g). TEV, Tobacco Etch Virus; MBP, maltose binding protein; SEC, size-exclusion chromatography

The dinucleotide c-di-GMP is rare in *Archaea* and *Eukarya* but ubiquitous in *Bacteria,* where it mediates a broad variety of environmental adaptations, including responses to O_2_, light, and nutrients ^8,10,11^. While the reaction of bacteria to changes in the O_2_ tension is often an inter- conversion between sessile and motile states, a switch between replicative and non-replicative states, and other equally far-reaching responses are possible ^12^. For example, the aerobic acetic acid bacterium *Gluconoacetobacter xylinus* expresses PDE-A1, an O_2_-regulated phosphodiesterase that limits the allosteric activation of the bacterial cellulose synthase to the air-water interface, where a cellulosic life raft gets made as the growth media become anaerobic ^10,13^. DosP is produced under control of the sigma factor RpoS exclusively during stationary phase and is then the most abundant *E. coli* c-di-GMP phosphodiesterase ^14^. The DosP O_2_ dose response is from ∼75 µM to atmospheric O_2_ (256 µM), consistent with the decline in O_2_ tension during that growth phase ^1^. The deletion of *dosP* from *E. coli,* is not associated with a gross defect in biofilm production but rather results in filamentous growth ^15^. In addition, an association of DosP with polynucleotide phosphorylase (PNPase) and an effect of c-di-GMP on that RNA-processing enzyme suggest a role in post-transcriptional regulation ^16^.

DosP occurs essentially unchanged in many enteric bacteria; it is also present in diverse bacterial species from other genera, including *Rhizobia, Agrobacteria,* and *Bordetella.* In *E. coli*, *dosP* is expressed from the *dosCP* operon together with *dosC*, which encodes an O_2_-sensing diguanylate cyclase^15^. Whereas O_2_-bound DosP degrades c-di-GMP to the linear dinucleotide pGpG ^1–3^; unliganded DosC protein (K_d(O2)_ of 20 µM) produces c-di-GMP from two molecules of GTP ^16^. We have hypothesized that this push-pull system establishes a rheostat mechanism for tight control of c-di-GMP levels within localized regions having PNPase ^16^. In support of such a role in post-transcriptional regulation, deletion of *dosC* (also called *dgcO)* from the plant pathogen *Pectobacterium carotovorum* was recently shown to impair its aerobic transcripts ^17^.

Structural characterization of DosP has been restricted to its isolated EAL and hPAS domains. The EAL domain crystallized as a dimer with a short helical loop in the substrate pocket instead of c-di-GMP ^18^. For the hPAS domain, crystal structures have been reported twice. In one report, the initially O_2_-bound domain crystallized as a mixed dimer with one monomer bound to O_2_, i.e. the oxy-state, and the other monomer in an unliganded state (PDB 1S66 and **Extended Data Fig. 1a**)^19^. In the other report, it was crystallized as a dimer in an oxidized state (PDB 1V9Y) and then reduced to the unliganded deoxy-state having Fe^2+^ heme (PDB 1V9Z)^20^. Kurokawa and colleagues have also deposited an unpublished structure with the hPAS dimer in a mixed state like PDB 1S66 (PDB 1VB6). To our knowledge, there is no available structure representing the oxy/oxy hPAS dimer or the c-di-GMP bound EAL-domain and therefore no useful structure-based models of the allosteric regulation.

Examinations of the hPAS domain have established that the heme is His-Fe-Met coordinated when unliganded ^19–21^. Therefore, for an exogenous ligand to bind to the iron atom, it must displace the internally-coordinated sulfur of a methionine, M95. Non-physiological ligands of ferrous heme like CO and NO can displace M95 to switch DosP to the ON-state^1,3^. An important difference of bound O_2_ from those ligands, however, is that O_2_ requires hydrogen bond donation from a universally conserved arginine residue of hPAS domains, R97 in DosP, for any significant binding to the iron atom ^19,22,23^. In DosP the competition of M95 against R97 for entry into the heme pocket is therefore presumed to be a key part of the switching mechanism^3,19^. Nevertheless, other than in the F-G loop, where M95 and R97 reside, only minimal differences have been observed between the mixed dimer structure of PDB 1S66 and the fully deoxy-state structure of PDB 1V9Z for the hPAS domain The structural model for the local conformation changes to the F-G loop that are induced by competition between M95 and R97 is summarized in **Extended Data Fig. 1b-d**..

Here, using cryo-electron microscopy (cryo-EM), we determined structures of full-length DosP in six states. These were the anaerobic state with substrate, the aerobic state with substrate, and an equilibrium of two states that occurred for each species (anaerobic and aerobic) when without substrate. Together with biochemical analyses, our results provide a structure-based framework to begin to understand how an allosteric signal is transmitted through DosP over more than 180 Å. Our work reveals that O_2_ regulates catalysis, not substrate binding. Conformation changes propagate from the regulatory hPAS domain to the EAL domain along connective helices and result in subtle changes to the position of catalytic metals in the active site.

## Results

### Oxy-DosP^WT^ and DosP^R97A^ offer stable ON and OFF states, respectively, for structural studies

To understand the architecture of DosP and the regulation of its phosphodiesterase by O_2_, we produced a maltose-binding protein (MBP)-tagged DosP by modifying a *malE-dosCP* construct that we previously described ^16^. We engineered a Tobacco Etch Virus (TEV) protease cleavage site before DosP such that the starting methionine would be the first residue of our scar-free, purified DosP (**Fig. 1b**). The MBP-DosP could be purified by amylose resin affinity and then size-exclusion chromatography (SEC). Subsequently the MBP tag was efficiently cleaved with TEV protease and then the DosP was separated from the MBP by a final SEC step (**Fig. 1c-d**). In air, the wild-type DosP (DosP^WT^) showed the expected oxy-state absorption, and under anaerobic conditions it showed a deoxy-state absorption; the spectra showed strong Soret bands and clear differentiation between the β- and ⍺-bands for each state (anaerobic maxima of β = 532 and ⍺ = 564 nm; aerobic maxima of β = 544 and ⍺ = 578 nm) (**Fig. 1e**). Because of the practical challenges to maintaining DosP^WT^ in the deoxy-state while preparing cryo-EM grids, we took advantage of an R97A substitution, which we previously showed weakens O_2_ binding by removing a stabilizing H-bond ^21^. DosP^R97A^ remained in the deoxy state even in air (**Fig. 1f**).

The phosphodiesterase activity was previously shown to be inhibited for deoxy-DosP^WT^ from measurements of the pGpG product ^1–3^. Here, we measured the disappearance of the c-di-GMP substrate by a modified commercial assay that uses a fluorescent reporter (see *Methods*). Under aerobic conditions, DosP^WT^ cleaved c-di-GMP, with an apparent k_cat_ of 3.2 s^-1^. By contrast, under anaerobic conditions, DosP^WT^ activity decreased to a k_cat_ of 0.4 s^-1^ (**Fig. 1g-h**). Consistent with the impaired O_2_ binding by DosP^R97A^, its phosphodiesterase activity under aerobic and anaerobic conditions was as low as that of anaerobic DosP (**Fig. 1g-h**). These results showed that DosP^WT^ was well behaved and switched ON by O_2_, whereas DosP^R97A^ remained in the OFF-state with or without O_2_.

### DosP adopts two global conformations in the absence of substrate

We prepared cryo-EM grids of DosP^WT^ without c-di-GMP under aerobic conditions and resolved two distinct conformations, which we termed “bent” and “straight” (**Extended Data Fig. 2-3, Table 1, and Fig. 2a-b**). The numbers of particles corresponding to each class were similar, with a relative ratio of 1.2 straight particle to 1 bent particle. The oxy-DosP^WT^ formed an elongated key-shaped dimer that resembled two seahorses joined by their tails. The dimeric status is consistent with size-exclusion chromatography multi-angle light-scattering experiments^24^. All the domains could be confidently docked with existing crystal structures or Alphafold 2 (AF2) predictions as the starting models. A striking result of this assembly was that roughly 180 Å separated the regulatory hPAS domain from the catalytic EAL domain (**Fig. 2a**). In each DosP^WT^ monomer the four structural domains followed the same order as in the linear sequence of amino acids. At the N-terminus was the hPAS domain. This was followed by a PAS-PAC, which consisted of a PAS domain without a cofactor but with a tightly integrated C-terminal PAC motif. Next came a silent cyclase domain with a canonical diguanylate cyclase structure but with EGTQF replacing the catalytic GGDEF motif. Finally, at the C-terminus, there was the EAL domain responsible for the phosphodiesterase activity (**Fig. 1a**, **Fig. 2a**). Those four structural domains were linked by well-resolved helices, consistent with a role in transmitting an allosteric signal from one end of the structure to the other. A notable exception to the well-resolved connective helices was a melted helix in one monomer of the bent form of DosP, discussed below. Together the paired domains and connective helices contributed an extensive dimer interface that totaled nearly 5,000 Å^2^.

**Figure 2.**
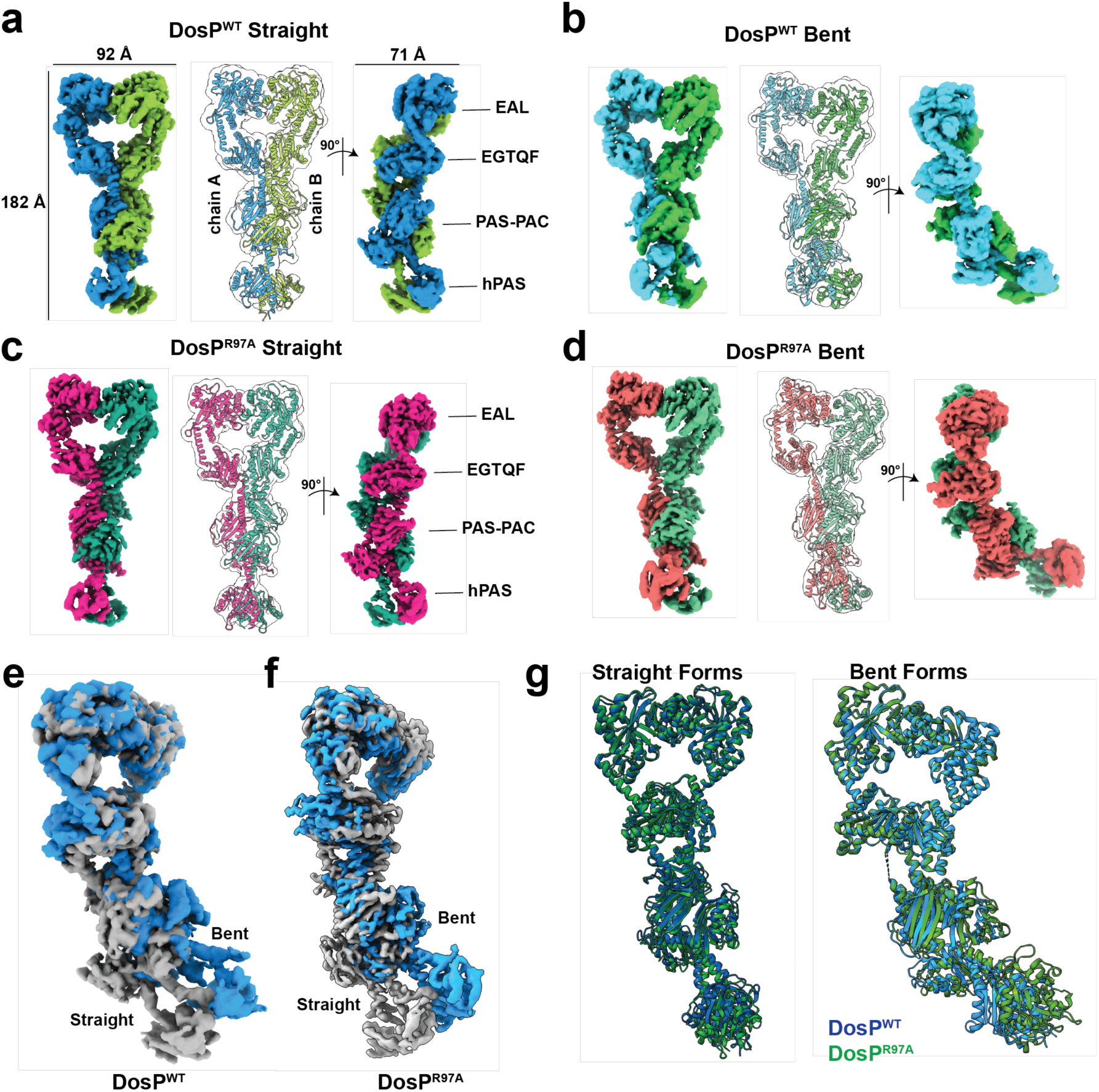
Structures of DosP^WT^ and DosP^R97A^. **a,** Cryo-EM map of the DosP^WT^ straight conformer. *Left:* map is shown at a contour of 0.0335 and colored by chain. Lines and labels indicate axial dimension. *Middle:* Atomic model depicted as cartoon and overlaid with a map that was gaussian filtered to level 1.5. *Right:* Map colored as in left panel and rotated 90°. Domains of DosP are labeled. **b,** Cryo-EM map of the DosP^WT^ bent conformer. *Left:* map is shown at a contour of 0.026 and colored by chain. *Middle:* Atomic model depicted as cartoon and overlaid with a map that was gaussian filtered to level 1.5. *Right:* Map colored as in left panel and rotated 90°. **c,** Cryo-EM map of the DosP^R97A^ straight conformer. *Left:* map is shown at a contour of 0.0142 and colored by chain. *Middle:* Atomic model depicted as cartoon and overlaid with a map that was gaussian filtered to level 1.5. *Right:* Map colored as in left panel and rotated 90°. Domains are labeled. **d,** Cryo-EM map of the DosP^R97A^ bent conformer. *Left:* map is shown at a contour of 0.075 and colored by chain. *Middle:* Atomic model depicted as cartoon and overlaid with a map that was gaussian filtered to level 1.5. *Right:* Map colored as in left panel and rotated 90°. **e,** DosP^WT^ bent form and straight form maps superposed through their EAL domains. The straight map is colored grey, and the bent form map is colored blue. **f,** DosP^R97A^ bent form and straight form maps superposed through their EAL domains. The straight map is colored grey, and the bent form map is colored blue. **g,** Atomic models of DosP^WT^ and DosP^R97A^ are superimposed. DosP^WT^ is depicted as blue cartoon and DosP^R97A^ is depicted as green cartoon. Bent and Straight forms are labeled. All panels are generated using ChimeraX. hPAS, heme-binding Per-Arnt-Sim; EAL, c-di-GMP phosphodiesterase domain; EGTQF-diguanylate cyclase domain.

**Table 1:** Cryo-EM data collection, refinement and validation statistics.

|  | #1 DosP<sup>FLAG</sup>+c-di-GMP (composite)<br>(EMDB-45746)<br>(PDB 9CMF) | #1 DosP<sup>FLAG</sup>+c-di-GMP (C-Terminal)<br>(EMDB- 45645) | #1 DosP<sup>FLAG</sup>+c-di-GMP (N-Terminal)<br>(EMDB-45646) | #1 DosP<sup>FLAG</sup>+c-di-GMP (Consensus)<br>(EMDB-45665) | #2 DosP<sup>R97A</sup> straight<br>(EMDB-44524)<br>(PDB 9BGV) | #3 DosP<sup>R97A</sup> bent<br>(EMDB-44646)<br>(PDB 9BKV) | #4 DosP straight<br>(EMDB-45485)<br>(PDB 9CDR) | #5 DosP bent<br>(EMDB-45489)<br>(PDB 9CE0) | DosP<sup>R97A</sup>+c-di-GMP (composite)<br>(EMDB-45727)<br>(PDB 9CLO) | DosP<sup>R97A</sup>+c-di-GMP (C-Terminal)<br>(EMDB-45667) | DosP<sup>R97A</sup>+c-di-GMP (N-Terminal)<br>(EMDB-45669) | DosP<sup>R97A</sup>+c-di-GMP (Consensus)<br>(EMDB-45670) |
| --- | --- | --- | --- | --- | --- | --- | --- | --- | --- | --- | --- | --- |
| **Data collection and processing** |  |  |  |  |  |  |  |  |  |  |  |  |
| Magnification | 81 kx | 81 kx | 81 kx | 81 kx | 81 kx | 81 kx | 105 kx | 105 kx | 165 kx | 165 kx | 165 kx | 165 kx |
| Voltage (kV) | 300 | 300 | 300 | 300 | 300 | 300 | 300 | 300 | 300 | 300 | 300 | 300 |
| Electron exposure (e-/Å<sup>2</sup>) | 50 | 50 | 50 | 50 | 50 | 50 | 50 | 50 | 50 | 50 | 50 | 50 |
| Defocus range (µm) | -0.8 to -2.0 | -0.8 to -2.0 | -0.8 to -2.0 | -0.8 to -2.0 | -0.8 to -2.0 | -0.8 to -2.0 | -0.6 to -1.6 | -0.6 to -1.6 | -0.8 to -2.0 | -0.8 to -2.0 | -0.8 to -2.0 | -0.8 to -2.0 |
| Pixel size (Å) | 0.5305 | 0.5305 | 0.5305 | 0.5305 | 0.5328 | 0.5328 | 0.4133 | 0.4133 |  |  |  |  |
| Symmetry imposed | C1 | C1 | C1 | C1 | C1 | C1 | C1 | C1 | C1 | C1 | C1 | C1 |
| Initial particle images (no.) | 9.8 million | 9.8 million | 9.8 million | 9.8 million | 10.5 million | 10.5 million | 5 million | 5 million | 2.5 million | 2.5 million | 2.5 million | 2.5 million |
| Final particle images (no.) | 307,809 | 307,809 | 157,857 | 157,857 | 353,471 | 391, 512 | 191,953 | 154,173 | 186,761 | 186,761 | 186,761 | 186,761 |
| Map resolution (Å) |  | 3.11 | 3.83 | 3.7 | 3.65 | 3.43 | 3.91 | 3.97 |  | 3.09 | 3.18 | 3.45 |
| FSC threshold |  | FSC = 0.143 | FSC = 0.143 | FSC = 0.143 | FSC = 0.143 | FSC = 0.143 | FSC = 0.143 | FSC = 0.143 |  | FSC = 0.143 | FSC = 0.143 | FSC = 0.143 |
| Map resolution range (Å) | 3.11-3.83 |  |  |  |  |  |  |  | 3.09-3.18 |  |  |  |
| **Refinement** |  |  |  |  |  |  |  |  |  |  |  |  |
| Initial model used (PDB code) | PDB 4HU4<br>PDB 1S66<br>AlphaFold 2<br>(Uniprot P76129) | - | - | - | DosP<sup>FLAG</sup> PDB 1S66<br>PDB 4HU4 | DosP<sup>FLAG</sup> PDB 1S66<br>PDB 4HU4 | DosP<sup>R97A</sup> PDB 1S66 | DosP<sup>R97A</sup> PDB 1S66 | DosP<sup>FLAG</sup> PDB 1S66<br>PDB 4HU4 | - | - | - |
| Model resolution (Å) | 3.6 | - | - | - | 3.9 | 3.8 | 7.2 | 6.9 | 3.1 | - | - | - |
| FSC threshold | FSC = 0.5 |  |  |  | FSC = 0.5 | FSC = 0.5 | FSC = 0.5 | FSC=0.5 | FSC=0.5 |  |  |  |
| Model resolution range (Å) |  |  |  |  |  |  |  |  |  |  |  |  |
| Map sharpening *B* factor (Å<sup>2</sup>) | deepenhancer | deepenhancer | deepenhancer | deepenhancer | deepenhancer | deepenhancer | deepenhancer | deepenhancer | deepenhancer | deepenhancer | deepenhancer | deepenhancer |
| **Model composition** |  |  |  |  |  |  |  |  |  |  |  |  |
| Non-hydrogen atoms | 12716 |  |  |  | 12590 | 12562 | 12604 | 12551 | 12680 |  |  |  |
| Protein residues | 1576 |  |  |  | 1574 | 1571 | 1574 | 1568 | 1574 |  |  |  |
| Ligands | HEM: 2<br>MG: 4<br>OXY: 2<br>C2E: 2 |  |  |  | HEM: 2 | HEM:2 | HEM: 2<br>OXY: 2 | HEM: 2<br>OXY: 2 | HEM: 2<br>MG: 4<br>C2E:2 |  |  |  |
| ***B* factors (Å<sup>2</sup>)** |  |  |  |  |  |  |  |  |  |  |  |  |
| Protein | 85.87 |  |  |  | 77.65 | 90.45 | 106.80 | 135.91 | 64.94 |  |  |  |
| Ligand | 77.88 |  |  |  | 133.97 | 119.08 | 168.94 | 227.53 | 62.49 |  |  |  |
| **R.m.s. deviations** |  |  |  |  |  |  |  |  |  |  |  |  |
| Bond lengths (Å) | 0.009 |  |  |  | 0.005 | 0.006 | 0.004 | 0.005 | 0.009 |  |  |  |
| Bond angles (°) | 0.801 |  |  |  | 0.692 | 0.839 | 0.983 | 1.001 | 1.095 |  |  |  |
| **Validation** |  |  |  |  |  |  |  |  |  |  |  |  |
| MolProbity score | 1.85 |  |  |  | 1.99 | 2.16 | 1.99 | 2.14 | 1.91 |  |  |  |
| Clashscore | 9.44 |  |  |  | 11.07 | 17.52 | 11.14 | 16.34 | 11.25 |  |  |  |
| Poor rotamers (%) | 0.00 |  |  |  | 0.00 | 0.44 | 0.00 | 0.15 | 0.00 |  |  |  |
| **Ramachandran plot** |  |  |  |  |  |  |  |  |  |  |  |  |
| Favored (%) | 94.91 |  |  |  | 93.50 | 93.74 | 93.44 | 93.60 | 95.03 |  |  |  |
| Allowed (%) | 5.09 |  |  |  | 6.50 | 6.13 | 6.37 | 6.27 | 4.90 |  |  |  |
| Disallowed (%) | 0.00 |  |  |  | 0.0 | 0.13 | 0.19 | 0.13 | 0.06 |  |  |  |

The moderate resolution of the oxy-DosP^WT^ map precluded confident modeling of the atomic structure. We considered the possibility that DosP^R97A^, which is locked in the deoxy state even in air, might offer higher resolution and provide insights into conformational changes associated with the switch from the ON to the OFF state. Therefore, we prepared grids with DosP^R97A^ and determined its structure using cryo-EM (**Sup. Fig. 3-4 and Table 1**). As hoped, DosP^R97A^ yielded a map with improved resolution and clear features (**Fig. 2 c-d**). Like oxy-DosP^WT^, DosP^R97A^ adopted straight and bent conformations for a similar number of particles in each class. The frequency of particles in the two states was similar, with 0.9 straight particle to 1 bent particle. In DosP^R97A^ the C-terminal regions were best resolved, with local resolution within the EAL domain approaching 3 Å (**Sup. Fig. 4c**). We modeled the hPAS dimer in DosP^R97A^ by docking two copies of the deoxy-state hPAS monomer from PDB 1S66 into the cryo-EM density. Although that region showed the lowest local resolution in our DosP^R97A^ map, the heme could be clearly visualized (**Sup. Fig. 5a**). A monomer EAL domain from PDB 4HU4 was also a close match to the cryo-EM map; however, the EAL-EAL dimer interaction from the crystal structure differed such that the N-terminus of one monomer shifted by ∼15Å when the crystal structure was superposed to a DosP^R97A^ EAL from the cryo-EM structure (**Extended Data Fig. 5b**).

**Fig. 2e** and **2f** present the cryo-EM maps of the straight and bent forms of DosP^WT^ and DosP^R97A^ aligned through their C-terminal EGTQF and EAL domains. In both proteins the N-terminal hPAS/PAS-PAC modules undergo a rigid body rotation of ∼30° between the two forms, straight vs. bent (**Fig. 2e-f**), but the N-terminal and C-terminal domains of DosP^WT^ and DosP^R97A^ align well when superposed separately (**Fig. 2g and Extended Data Fig. 5c**). A previous crystal structure of the isolated EAL domain of DosP without substrate showed a helix comprised of residues 643-648 projecting into the putative substrate-binding pocket, which would prevent c-di-GMP binding (**Extended Data Fig. 5d**). This loop was previously termed an “inhibitory loop;” however, the loop does not appear functionally to inhibit substrate binding (discussed below). Therefore, we renamed it the “q-loop” because of the shape it adopts when it partly occupies the substrate-binding site. In the absence of c-di-GMP, the straight and bent structures for DosP^WT^ and DosP^R97A^ showed the q-loop to be in the same position, suggesting its placement was not regulated by O_2_. The limited resolution did not allow us to infer detailed changes between the DosP^WT^ and DosP^R97A^ structures without substrate, other than to note that in the bent forms of the two structures the hPAS domains were positioned slightly differently when the structures were aligned to their EAL domains (**Fig. 2g**).

We analyzed the bent vs straight forms from the maps of DosP^R97A^ because of their higher resolution. The nexus of the change between straight and bent was a “hinge helix,” consisting of residues 352-382, which connected the PAS-PAC to the EGTQF domain in both monomers of the straight form. In the bent conformation, the hinge helix was melted in one monomer, rendering residues 379-383 of that monomer unresolvable (**Fig. 3a**). Farther along the structure, within the EGTQF and EAL domains, the bent form of DosP^R97A^ adopted a slightly wider conformation, which was most obvious for the EGTQF-EAL connector helix (residues 539-562) (**Fig. 3b**). Those changes propagated from the EGTQF dimer where a loop, comprising residues 384-395, rearranged to create a more extensive dimer interface in the bent form, compared to the straight form (**Fig. 3c**). The 384-395 loop comes directly after the hinge helix, revealing a coupling of the melted helix to the conformation changes in the C-terminal EGTQF and EAL domains.

**Figure 3.**
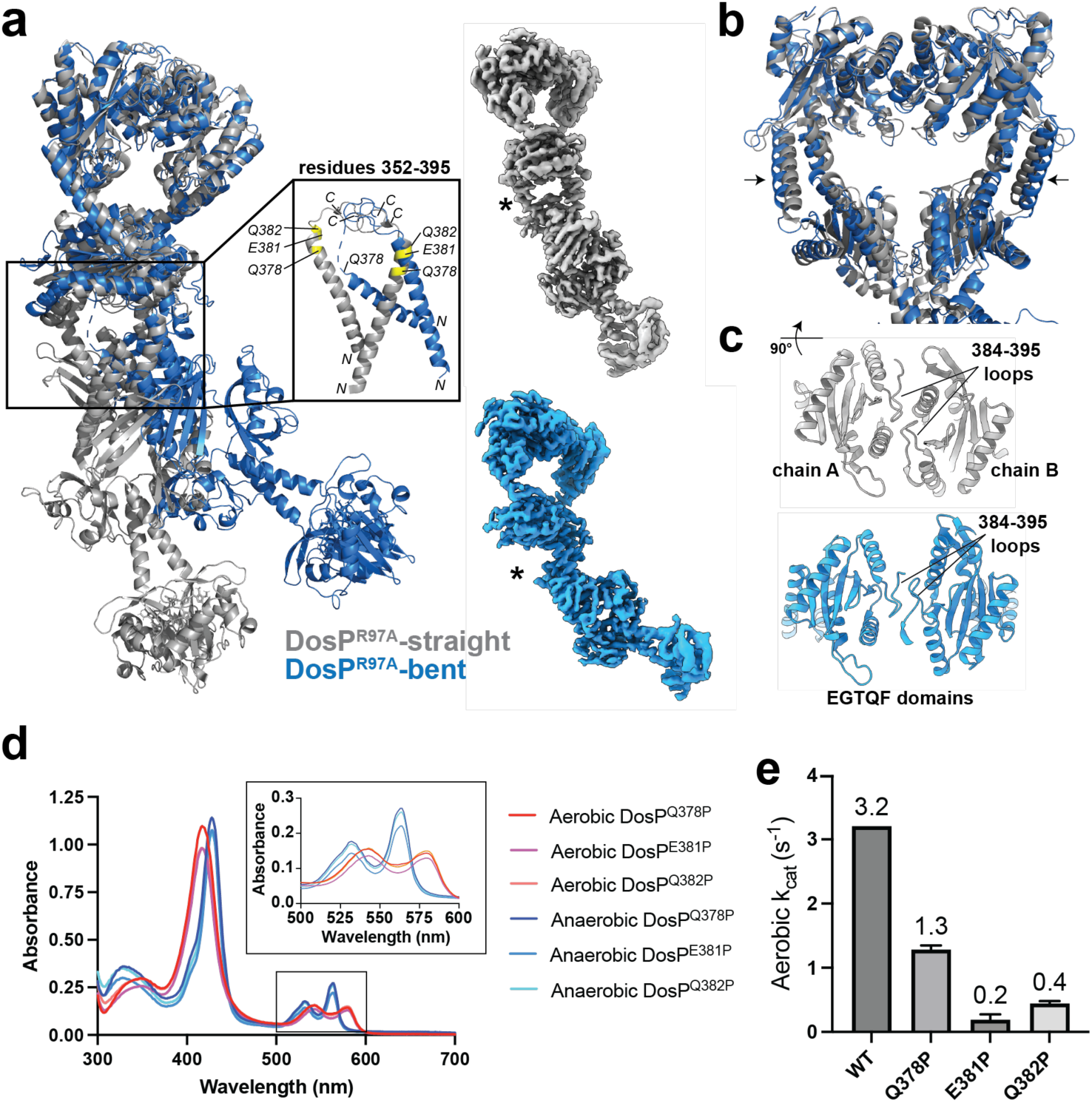
Investigation of the PAS-PAC/EGTQF hinge of DosP. **a,** Atomic models of the straight and bent forms of DosP^R97A^ depicted as grey cartoon or blue cartoon, respectively. Box highlights the PAS-PAC/EGTQF hinge helix (residues 352-282). Middle inset: Atomic models of residues 352-395 of the bent and straight forms of DosP^R97A^. NH_2_ and COOH termini are labeled with *N* and *C*, respectively. Positions of residues interrogated by mutagenesis are labeled yellow on the model of DosP^R97A^ straight. Top right inset: Cryo-EM map of DosP^R97A^ straight form. Asterisk denotes the well-resolved hinge helix. Bottom right inset: Cryo-EM map of DosP^R97A^ bent form. Asterisk denotes the unresolved hinge helix. **b,** Atomic models of the EAL and EGTQF domains of the straight and bent forms of DosP^R97A^ depicted as grey cartoon or blue cartoon, respectively. Arrows indicate changes of the helix between the EAL and EGTQF domains. **c,** The EGTQF dimer interface, viewed as if the position in (b) was rotated 90° upwards towards the reader. Top: the straight form of DosP^R97A^ shown as grey cartoon. The two chains and dimer interface loops are labeled. Bottom: the bent form of DosP^R97A^ shown as blue cartoon. **d,** Absorption spectra of DosP^Q378P^, DosP^E381P^, and DosP^Q382P^ under aerobic conditions (red, magenta, and orange, respectively), compared to anaerobic conditions (dark blue, blue, and cyan, respectively). **e,** Bar graph showing the k_cat_ values determined from the turnover of 2 µM c-di-GMP at 25°C by 10 nM aerobic: DosP^WT^, DosP^Q378P^, DosP^E381P^, and DosP^Q382P^. In (d) and (e) the assay buffer was 20 mM Tris-HCl, 2.5 mM MgCl_2_, 2.0 mM DTT, pH 8.0. Cartoons depicted in (a) and (b) generated using Pymol. Inset maps in (a) and cartoons in (c) generated using ChimeraX. EAL, c-di-GMP phosphodiesterase domain; EGTQF-diguanylate cyclase domain.

### Critical role of the PAS-PAC/EGTQF hinge helix in regulating the DosP phosphodiesterase

We tested whether the hinge helix between the PAS-PAC and EGTQF domains contributed to the regulation of DosP by O_2_. We reasoned that breaking this helix with a proline substitution would prevent DosP from adopting the straight conformation. To this end, we selected for interrogation residues Q378, E381, and Q382 from that region and generated independent substitutions of those residues with proline. The mutated proteins were recovered in yields comparable to wild type DosP and eluted at identical volumes by SEC (**Extended Data Fig 5e**); in addition, they showed excellent oxy-state and deoxy-state absorption spectra under aerobic and anaerobic conditions, respectively (**Fig. 3d**) Under aerobic conditions, however, DosP^Q378P^, DosP^E381P^, and DosP^Q382P^ showed reduced activity compared to DosP^WT^ (**Fig. 3e**). The observation of deoxy-state behavior in O_2_-ligated proteins with intact phosphodiesterase domains supports a role for the intervening regions in allosteric communication between the hPAS and EAL domains.

### Structural characterization of substrate-bound DosP in the ON-state

Because we did not observe a unique overall conformation when comparing DosP^R97A^, which was locked in the inactive deoxy-state, to oxy-DosP, which was active, we hypothesized that a switch to the ON-state might require the coordinated associations of O_2_ and c-di-GMP. For the examinations of substrate-bound DosP we utilized, for convenience, a full-length DosP harboring three copies of the FLAG motif on its C-terminus (hereafter DosP^FLAG-WT^). DosP^FLAG-WT^ was easily purified to homogeneity by anti-FLAG affinity resin and then SEC, from which it migrated as a single peak (**Extended Data Fig. 6a**). The C-terminal epitope tag had no effect on heme binding, O_2_ binding, or regulated enzymatic activity (**Extended Data Fig. 6b-c**). Isothermal titration calorimetry (ITC) of DosP^FLAG-WT^ binding to c-di-GMP in the presence of inhibitory calcium revealed a strong interaction with an apparent affinity equal to or less than 10 nM (**Extended Data Fig. 6d**). Owing to the small heat signatures, a more precise measurement of the affinity could not be made; however, based on these ITC measurements, the stoichiometry of the interaction was one molecule of c-di-GMP per monomer of DosP^FLAG-WT^, suggesting that the EAL domain was the only binding site for the c-di-GMP. The affinity of DosP^FLAG-WT^ for c-di-GMP was stronger than the K_D_ of ∼120 nM measured by ITC for the non-catalytic EAL domain of FimX ^25,26^ or the K_D_ of ∼2 µM measured by ITC for the interaction between c-di-GMP and an EAL domain from *Vibrio cholerae* ^27^.

For structural studies, we purified the DosP^FLAG-WT^ in air and supplemented it with catalytic magnesium (MgCl_2_) after SEC and added substrate (c-di-GMP) seconds before freezing the cryo-EM grids. In striking contrast to the protein examined without substrate, the substrate-bound DosP^FLAG-WT^ adopted a single global conformation (**Fig. 4a, Extended Data Fig. 3, Extended Data Fig. 6e, and Extended Data Fig. 7**). Extensive 3D classification efforts failed to identify additional conformations. By comparison, the bent and straight species without substrate readily emerged in the *ab initio* map generation stages of data processing (**Extended Data Figs. 2c and 4c**). Local refinement of the hPAS/PAS-PAC domains or the EGTQF/EAL domains generated maps with improved features (**Extended Data Fig. 7c-f**). The maps superposed well, and we generated a single composite map for illustrations and model building (**Extended Data Fig. 7h-i**). Substrate-bound DosP^FLAG-WT^ resembled the straight form of DosP^WT^ or DosP^R97A^ (**Extended Data Fig. 6e**). In each monomer of the dimeric DosP^FLAG-WT^, the EAL domain contained an intact c-di-GMP that was well-resolved in the map (**Fig. 4c-d**). A rearrangement of q-loop of DosP^FLAG-WT^ moved the L647 residue ∼8.1 Å out of the binding pocket to make room for the c-di-GMP, which had clashed with L647 in the substrate-free structures (**Fig. 4e**).

**Figure 4:**
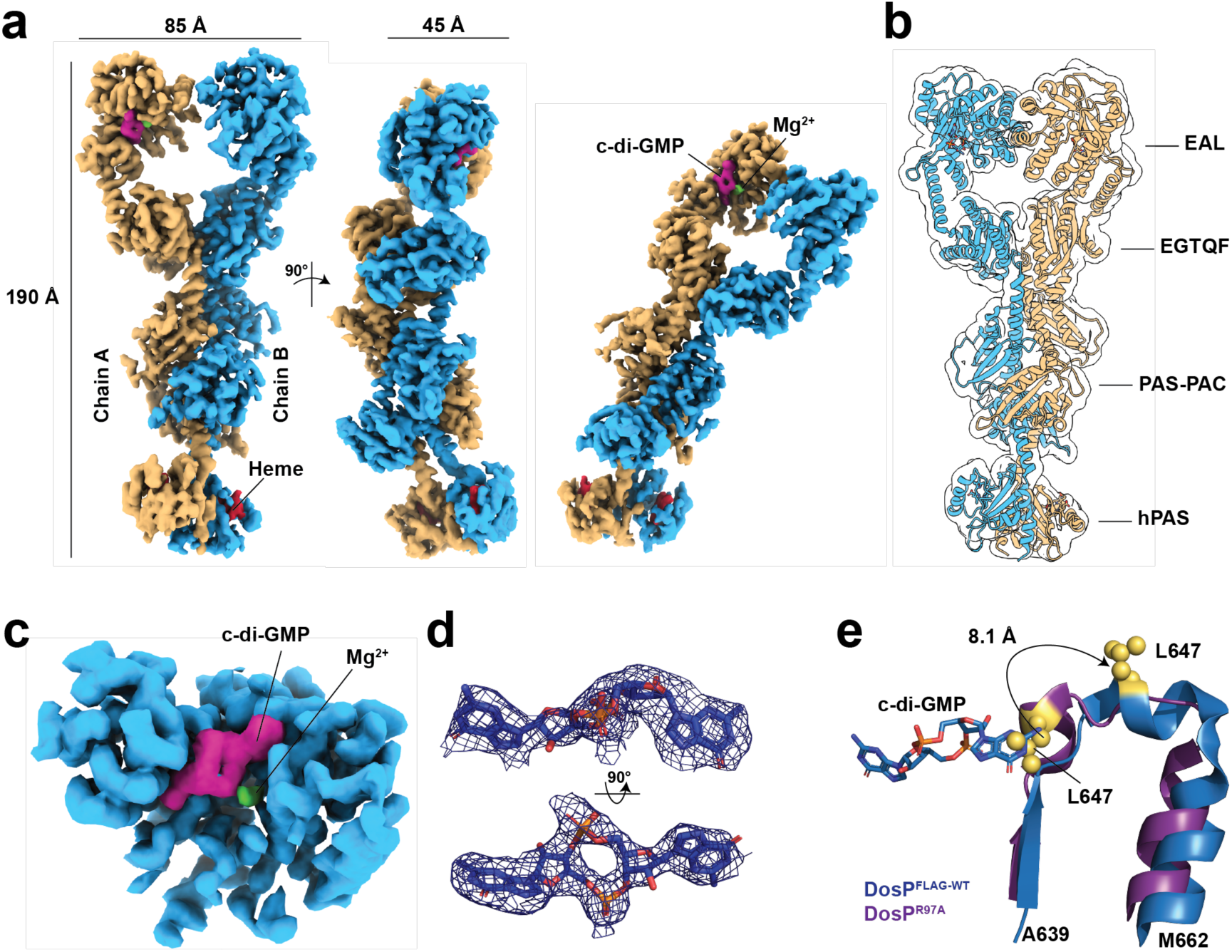
Structural details of DosP^WT^ with bound c-di-GMP. **a,** Composite map of DosP^FLAG-WT^ with the bound c-di-GMP from three orientations. The left panel displays the two chains as cyan and yellow density maps at a contour level of 0.064. The density for c-di-GMP and heme is shown in magenta and red, respectively. The density of magnesium is shown in green. Axial dimensions are labeled. **b,** Atomic model depicted as cartoon and overlaid with a map that was gaussian filtered to level 1.5. Domains of DosP are labeled. **c,** close view of a single EAL domain with bound c-di-GMP. Density for the EAL domain is colored cyan, c-di-GMP density is colored magenta, and magnesium density is colored green. **d,** Density for c-di-GMP shown in two orientations. The c-di-GMP is represented as sticks and the density map displayed as a blue mesh carved at 2.0 Å and sigma=10. **e,** DosP^FLAG-WT^ and DosP^R97A^ residues 639-662 superposed and displayed with blue and purple cartoon, respectively. L647 is depicted with yellow spheres and c-di-GMP is depicted in sticks. Arrow indicates the change in position of L647 between the two forms of DosP. Panels a-c generated using ChimeraX. Panels d-e generated using Pymol. hPAS, heme-binding Per-Arnt-Sim; EAL, c-di-GMP phosphodiesterase domain; EGTQF-diguanylate cyclase domain.

As a rule, EAL domains coordinate two magnesium atoms (Me1 and Me2) to activate a catalytic water molecule (W1). A conserved lysine residue assists the deprotonation of W1 to generate a nucleophilic hydroxyl that attacks the scissile phosphate of c-di-GMP; this lysine is K726 in DosP. A second water molecule (W2) may serve as a proton donor for the leaving O3’ group ^27,28^. Although the resolution of our reconstruction precluded modeling the water molecules, the catalytic magnesiums could be clearly visualized and assigned by reference to the crystal structure for TBD1265 EAL (PDB 3N3T) (**Fig. 4c and Fig. 5a**). The metals were situated on an acidic surface that included E584 of the namesake EAL motif. Consistent with previous characterizations of EAL domains with substrate, the c-di-GMP was in an extended conformation ^27,28^. The non-scissile phosphate did not interact with either metal atom but formed an electrostatic interaction with the guanidinium side chain of R588 (**Fig. 5b**). The two guanine bases, G1 and G2, were differently recognized: G1 was largely buried and held by non-specific hydrophobic interactions; G2 was more accessible to solvent but engaged in more specific interactions, including a pi-stacking interaction with Y784. To test the contributions of those substrate-pocket residues to the enzymatic activity, we determined the rates of DosP^E584A^, DosP^R588A^, DosP^R588E^, and DosP^Y784A^ turnover of c-di-GMP. As anticipated, DosP^E584A^ was entirely inactive (**Fig. 5c**). Both DosP^R588A^ and DosP^Y784A^ showed reduced aerobic activity, consistent with those residues’ roles in substrate binding. The charge-swapped DosP^R588E^ substitution resulted in a complete lack of aerobic activity in accord with that residue’s electrostatic interaction with the non-scissile phosphate of c-di-GMP (**Fig. 5c**). Altogether, these results indicate that DosP possesses a canonical EAL substrate binding pocket.

**Figure 5:**
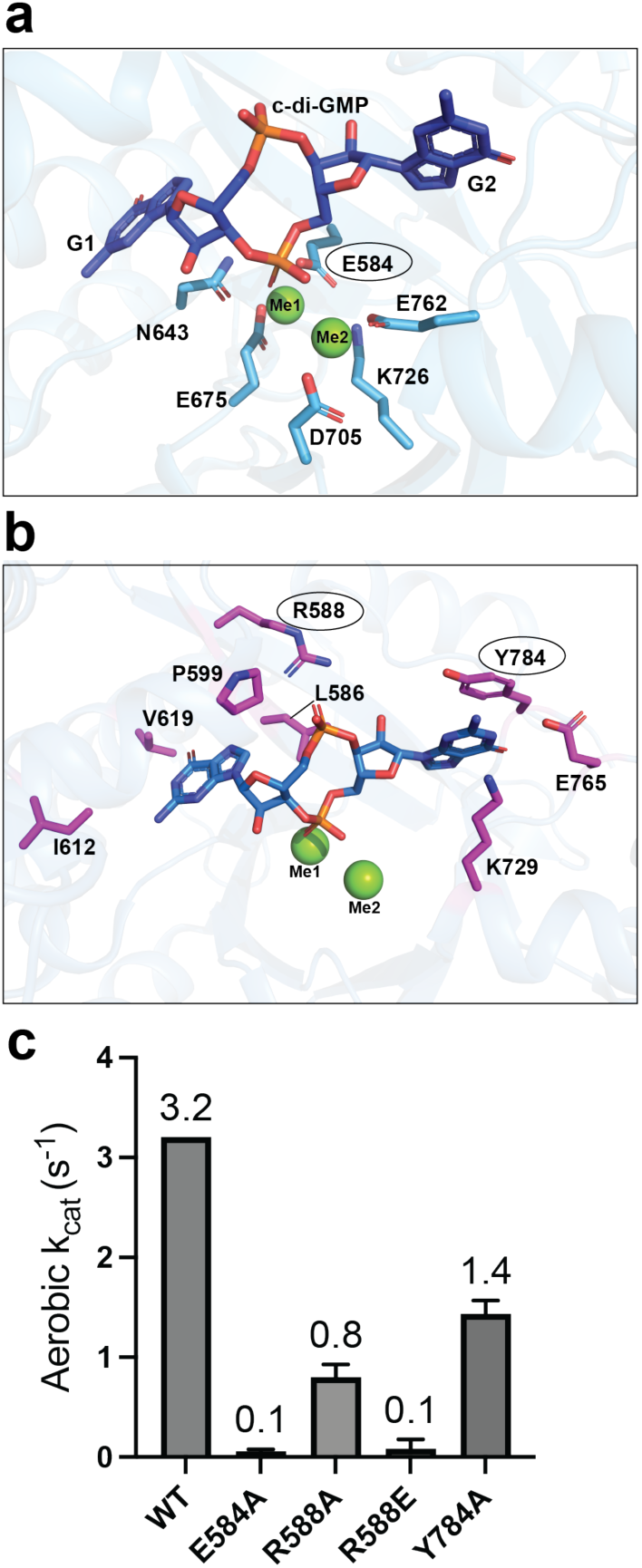
The DosP catalytic pocket. **a,** Bound c-di-GMP is shown as blue sticks. The guanine bases are labeled G1 and G2. The two magnesium ions are shown as green spheres and labeled Me1 and Me2. Metal-coordinating residues are depicted as cyan sticks. The rest of the EAL domain is depicted as a transparent cartoon. **b,** Residues interacting with c-di-GMP are shown as magenta sticks. The rest of the EAL domain is depicted as transparent cartoon. **c,** Bar graph showing the k_cat_ values determined from the turnover of 2 µM c-di-GMP at 25°C by 10 nM aerobic DosP^WT^, DosP^E584A^, DosP^R588A^, DosP^R588E^, and DosP^Y784A^. The buffer for the assays was 20 mM Tris-HCl, 2.5 mM MgCl_2_, 2.0 mM DTT, pH 8.0. Residues whose substitutions were tested in (c) are highlighted with circles in (a) and (b). Panels a and b generated with PyMOL.

### Regulatory conformational changes between substrate-bound ON vs. OFF states

Based on our experience determining structures of DosP without substrate, we reasoned that we might obtain a map of similar or better quality with DosP^FLAG-R97A^ bound to c-di-GMP. However, it was plausible to consider that the anaerobic state might inhibit substrate binding. To test this, we carried out a simple binding assay based on the observation that c-di-GMP (MW 690.1) readily passed through a centrifugal concentrator with a molecular weight cut-off of 100,000. When equal concentrations of DosP and c-di-GMP were incubated in the presence of inhibitory calcium, the binding of c-di-GMP to DosP prevented its passage through the filter. Aerobic and anaerobic DosP retained essentially all the c-di-GMP at concentrations of 125 nM and 250 nM (**Extended Data Fig. 8a**). These data show that any modest change in affinity for c-di-GMP beyond the sensitivity of this assay cannot explain the 8-fold difference in activity in conditions with 2 µM c-di-GMP (**Fig. 1g**). To test this hypothesis more fully, we incubated DosP^WT^and DosP^R97A^ with 100 µM c-di-GMP and assayed their phosphodiesterase activities. Despite the 50-fold increase in substrate concentration, DosP^R97A^ remained unstimulated, with k_cat_ values identical to what we observed in the typical experiment with 2 µM substrate (**Fig. 6a**). This led us to consider the possibility that O_2_ ligation regulates catalysis but not substrate binding. We therefore collected cryo-EM data with DosP^FLAG-R97A^ bound to c-di-GMP. As hoped, this sample yielded our best map, with good density throughout the protein and clear density for the bound substrate (**Extended Data. 3, Extended Data Fig. 8b-f and Fig. 6b-e**). Like DosP^FLAG-WT^, substrate-bound DosP^FLAG-R97A^ adopted a single conformation that largely resembled the straight form (see below). The c-di-GMP was well-resolved, and the q-loop was displaced (**Fig. 6e**). Taken together, these data suggest that the OFF-state does not abolish substrate binding but instead impairs a downstream, catalytic step.

**Figure 6:**
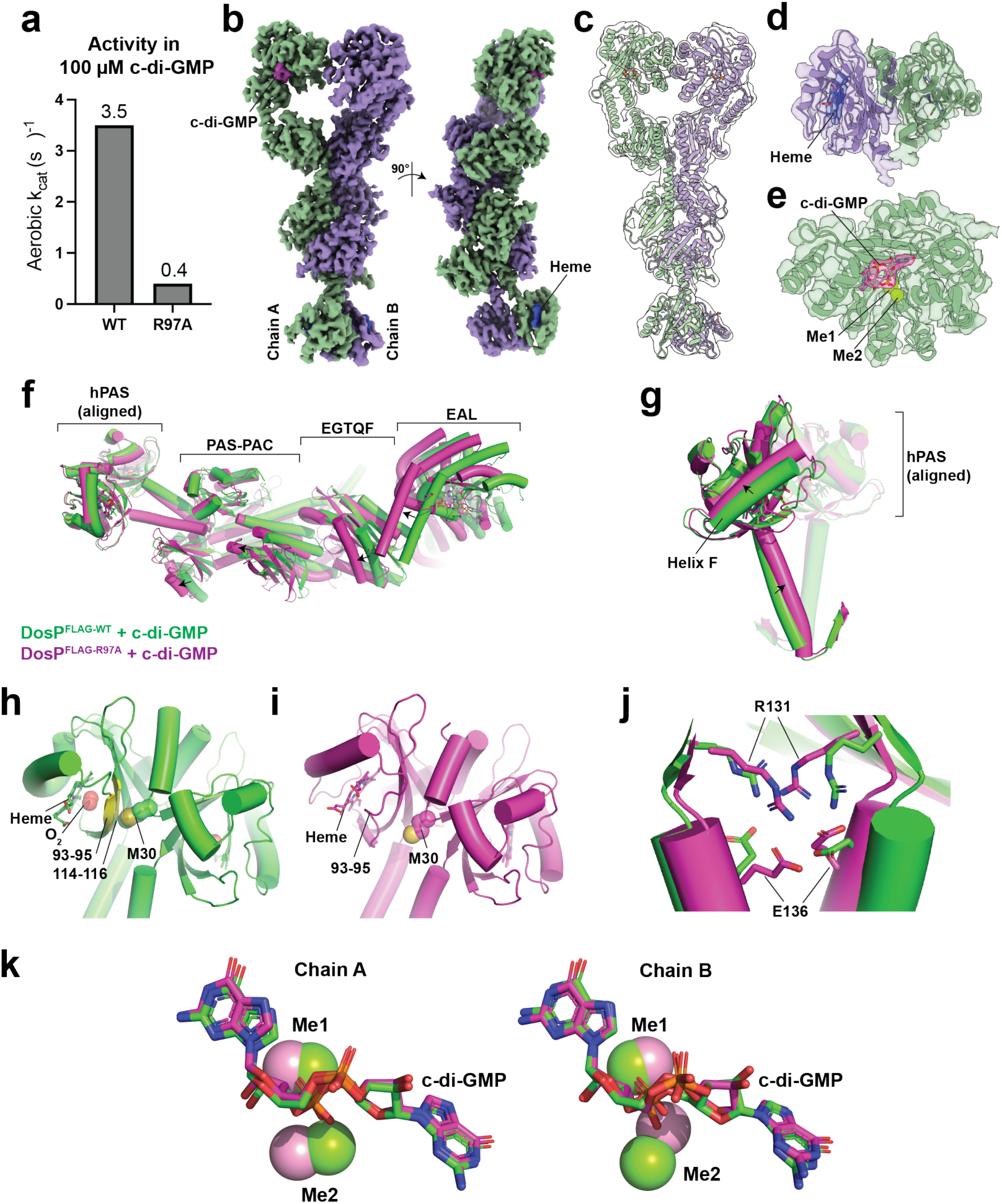
Structural details of DosP^R97A^ with bound c-di-GMP. **a,** Turnover of 100 µM c-di-GMP at 25°C by 10 nM aerobic DosP^WT^, and DosP^R97A^. The buffer for the assays was 20 mM Tris-HCl, 2.5 mM MgCl_2_, 2.0 mM DTT, pH 8.0. **b,** Cryo-EM map of substrate-bound DosP^FLAG-R97A^ in complex with c-di-GMP that is colored magenta and with the two chains colored as mint green and lavender. **c,** Atomic model depicted as cartoon and overlaid with a map that was gaussian filtered to level 1.5. **d,** Density and atomic model for the hPAS domains with their bound heme. **e,** Density and atomic model for one of the EAL domains with its bound c-di-GMP and magnesium ions. **f,** Atomic models of substrate-bound DosP^FLAG-WT^ and ^DosPFLAG-R97A^ aligned by their hPAS domains and colored green and magenta, respectively. Domains are labeled with brackets. Arrows indicate differences in conformation. **g,** The hPAS domains of substrate-bound DosP^FLAG-WT^ and ^DosPFLAG-R97A^ aligned and colored as in (f). One monomer is set partially transparent. Differences in conformation and Helix F are labeled. **h,** The hPAS dimer of DosP^FLAG-WT^ depicted as green cartoon. Heme depicted as sticks. Residues 93-95 and 114-114 of one monomer are highlighted as yellow cartoon and the side chain atoms of M30 from the other monomer are depicted with spheres. **i,** The hPAS dimer of DosP^FLAG-R97A^ depicted as magenta cartoon. Heme depicted as sticks. Residues 93-95 of one monomer are highlighted as yellow cartoon and the side chain atoms of M30 from the other monomer are depicted with spheres. **j,** A hydrogen bond between R131 and E136 links the hPAS domain to the helix leading to the PAS-PAC domain. DosP^FLAG-WT^ and ^DosPFLAG-R97A^ aligned through their hPAS domains are depicted as green and magenta cartoons, respectively. R131 and E136 are depicted as sticks. **k,** changes in the position of Me1 and Me2 between DosP^FLAG-WT^ and DosP^FLAG-R97A^. The EAL domains of each chain were superposed. The c-di-GMP is shown as sticks and colored by heteroatom and species of DosP as in (f). The magnesium ions shown as green or pink spheres correspond to DosP^FLAG-WT^ and DosP^FLAG-R97A^, respectively. Panels b-e generated using ChimeraX. Panels f-k generated using Pymol. hPAS, heme-binding Per-Arnt-Sim; EAL, c-di-GMP phosphodiesterase domain; EGTQF-diguanylate cyclase domain.

Our cryo-EM structures of the substrate-bound oxy/oxy ON-state structure (DosP^FLAG-WT^)showed clear differences from the deoxy/deoxy OFF-state structure (DosP^FLAG-R97A^). Superposing the substrate-bound structures of DosP^FLAG-WT^ and DosP^FLAG-R97A^ through their hPAS domains revealed the DosP^FLAG-R97A^ to be in a compressed form (**Fig. 6f**). The sites of those differences included the F-helix with the H77 residue of proximal iron coordination, the F-G loop with M95 and R97, and the linker helix from the hPAS to the PAC-PAS domain (**Fig. 6g**). The differences between the hPAS dimers of the oxy/oxy DosP^FLAG-WT^ and the deoxy/deoxy DosP^FLAG-R97A^ contrasts with the crystal structures of the mixed-state (PDB 1S66) and the fully deoxy state (PDB 1V9Z, see **Extended Data Fig. 1d**). Examining the hPAS dimer interface in the ON-state, i.e. in substrate-bound oxy-DosP^FLAG-WT^, we noticed that one monomer’s M30 residue contacted residues 93-95 in βG and 114-116 in βH of the other monomer (**Fig. 6h**). This interaction appeared to stabilize the F-G loop’s O_2_-bound conformation. By contrast, in the OFF-state conformation, i.e. in substrate-bound DosP^FLAG-R97A^, the M30 residue could not contact the other monomer because, upon coordination of the M95 sulfur to the heme iron, the F-G loop moved away from M30 (**Fig. 6i**). The stabilization of the F-G loop either by M30 from the other monomer in the ON-state, or else by the ligation of heme iron by M95 in the OFF-state, is apparent in the cryo-EM density for these domains. In the ON-state, F-G loop residues 87-92 are not resolved, but density becomes apparent at residue 93 (**Extended Data Fig. 9a**). In the OFF-state, the full F-G loop is resolved (**Extended Data Fig. 9b**). The poor density for the F-G loop in the oxy-state is unlikely to be entirely due to the lower resolution, as the highest B-factors of the 1.8 Å crystal structure of the mixed hPAS dimer (PDB 1S66) belong to residues 89-91 of the oxy-state F-G loop, and those residues were modeled with an occupancy of 0^19^. The hPAS domain connects to the PAS-PAC domain with R131 of the βI strand forming a hydrogen bond with E136 in the connective helix (**Fig. 6j**). We hypothesized that this interaction may facilitate the response of that helix to the regulatory changes in the hPAS.

To gain insights into regulatory change in the catalytic domain, we superposed the EAL domains of ON-state substrate-bound DosP^FLAG-WT^ and the OFF-state DosP ^FLAG-R97A^. The location of the catalytic magnesium metals differed between these two forms, with Me2 showing the clearest differences (**Fig. 6k**). Though the modest resolution does not allow confident modeling of the side-chain movements governing the changed position of magnesium ions, it is reasonable to suppose that a shift of Me2 could underlie the switch to the OFF-state in anaerobic DosP (**Extended Data Fig. 9c-d**).

To investigate the communication to the EAL domain from the hPAS and PAC-PAS domains, we prepared samples of DosP harboring alanine substitutions at M30 and R131 and examined their ability to bind O_2_ and regulate the phosphodiesterase. DosP^M30A^ showed a reduced affinity for O_2_ that was apparent from the similarity of the aerobic absorption spectra to the spectra obtained under anaerobic conditions (**Fig. 7a**). Consistent with its lower affinity for O_2_, DosP^M30A^showed only weak phosphodiesterase activity that could not be stimulated by air (**Fig. 7b** note, DosP^WT^ data from Fig. 1 is plotted for comparison). Additionally, we saturated DosP^M30A^ with CO, a heme ligand that is not physiological but that can activate the wild type protein. However, the CO-bound DosP^M30A^ was inactive despite presenting a clear carbonmonoxy absorption spectrum (**Fig. 7b and Extended Data Fig. 9e and 1**). These results are consistent with the M30 side chain normally serving to stabilize the ON-state.

**Figure 7.**
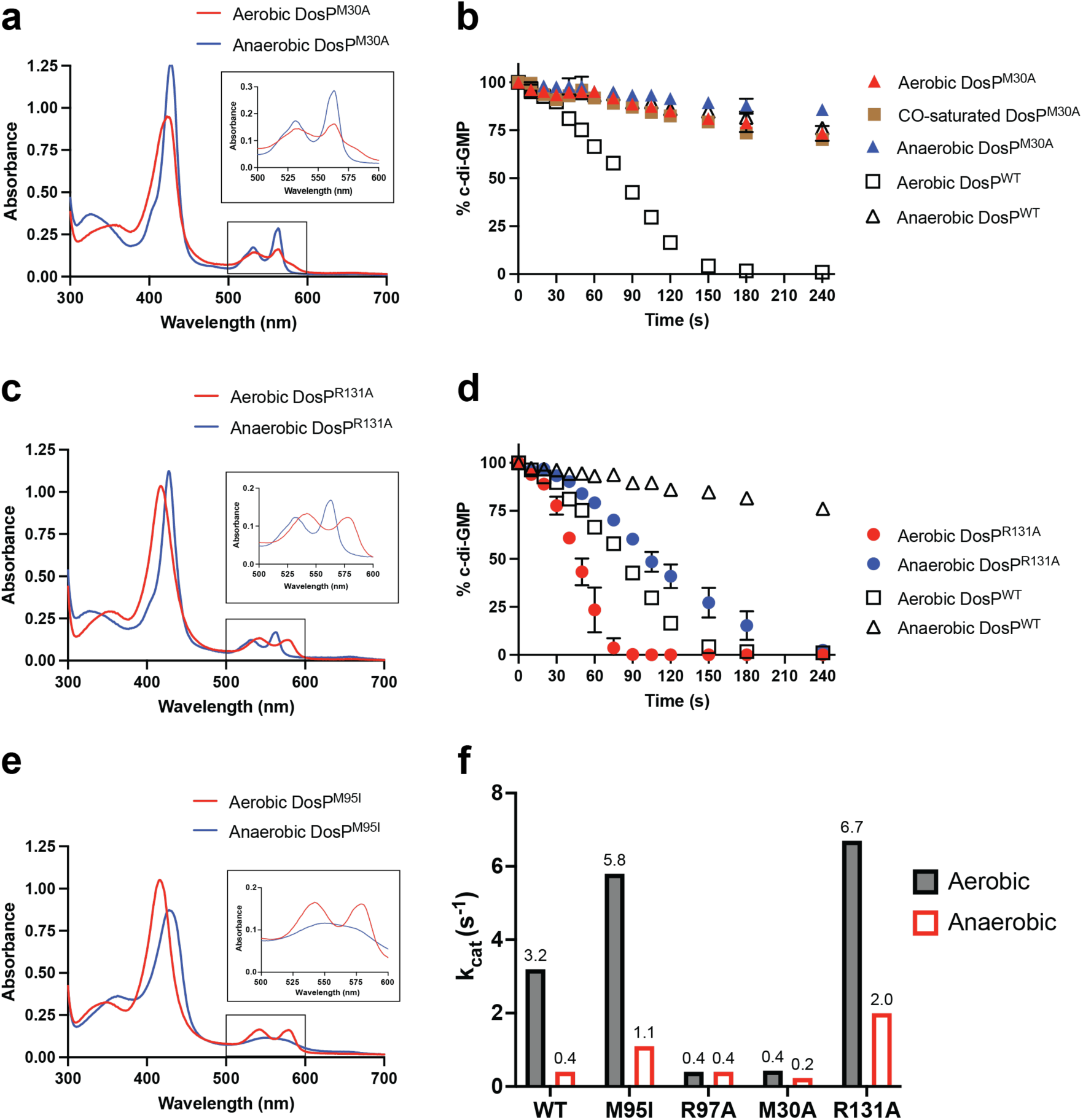
Effect of allosteric hPAS domain mutations. **a,** Absorption spectra of DosP^M30A^ under aerobic conditions (red), compared to anaerobic conditions (blue). **b,** Turnover of 2 µM c-di-GMP at 25°C by 10 nM aerobic DosP^M30A^ (red triangles), compared to anaerobic DosP^M30A^ (blue triangles) and CO-saturated DosP^M30A^ (orange squares). **c,** Absorption spectra of DosP^R131A^ under aerobic conditions (red), compared to anaerobic conditions (blue). **d,** Turnover of 2 µM c-di-GMP at 25°C by 10 nM aerobic DosP^R131A^ (red circles), compared to anaerobic DosP^R131A^ (blue circles). For comparison, the enzymatic activities of aerobic DosP^WT^ (open squares) and anaerobic DosP^WT^ (open triangles) are re-plotted from Fig. 1g in (b) and (d). **e,** absorption spectra of DosP^M95I^ under aerobic conditions (red), compared to anaerobic conditions (blue). **f,** Bar graph showing the k_cat_ values determined from the turnover of 2 µM c-di-GMP at 25°C by 10 nM aerobic DosP^WT^, DosP^M95I^, DosP^R97A^, DosP^M30A^, and DosP^R131A^. The assay buffer was 20 mM Tris-HCl, 2.5 mM MgCl_2_, 2.0 mM DTT, pH 8.0.

In contrast to M30, the R131 residue did not appear to contribute to O_2_ binding, and a DosP^R131A^ mutant bound O_2_ comparably to DosP^WT^ (**Fig. 7c**). Nevertheless, an R131A substitution was associated with a marked uncoupling of the hPAS from the phosphodiesterase: specifically, deoxy-DosP^R131A^ was not inhibited (k_cat_ = 2.0 s^-1^), and oxy-DosP^R131A^ was more than twice as active (k_cat_ = 6.7 s^-1^) as oxy-DosP^WT^ (**Fig. 7d**). As a final comparison, we generated DosP^M95I^, a mutation that raised the affinity for O_2_ about 10-fold by eliminating the competition from the endogenous M95 sulfur for the heme iron ^21^. Anaerobic DosP^M95I^ was readily distinguishable from the other versions of anaerobic DosP in that it produced an absorption spectrum for the deoxy state that corresponded to a pentacoordinate ferrous heme iron (**Fig. 7e**). Compared to the inactive deoxy-DosP^WT^, deoxy-DosP^M95I^ was active (k_cat_ = 1.1 s^-1^) and the DosP^M95I^ aerobic activity (k_cat_ = 5.8 s^-1^) was nearly twice that of DosP^WT^ (**Fig. 7f**).

## Discussion

The data herein provide a structural framework to understand the large family of heme-based oxygen sensor-effector proteins. DosP is both a model O_2_ sensor and prototypical c-di-GMP phosphodiesterase for study. Its environmental signal, O_2_, is known, and the ligation of this signal can be quantified. Therefore, DosP offered an opportunity to examine the regulatory conformational changes in a multi-domain, signal-transducing enzyme. We used cryo-EM and biochemical methods to show that DosP is an elongated dimer with the regulatory hPAS domain separated from the catalytic phosphodiesterase domain by ∼180 Å. Our study took advantage of the fast time scale of cryo-EM grid freezing to capture c-di-GMP bound to the EAL domain with catalytic magnesium; and our structural analysis supports the two-metal reaction mechanism proposed by earlier studies^11,27–30^. The elongated nature of the DosP dimer observed for both the oxy-state (ON) and deoxy-state (OFF) implies that the allosteric regulation travels linearly from the hPAS domain to the EAL domain (**Fig. 8a**). The linkage of the DosP domains by single-helix connectors enabled structure-guided mutations that decoupled the hPAS domain from the EAL domain but left intact the ability of the hPAS domain to bind O_2_. Experiments with DosP^R131A^ showed that decoupling the hPAS domain from the EAL domain can result in phosphodiesterase activity for the normally OFF-state deoxy protein. These findings are consistent with reports of activity for the isolated EAL domain^18^ as well as for DosP^H77A^, where mutation of the proximal histidine abolished heme binding^31^.

**Figure 8.**
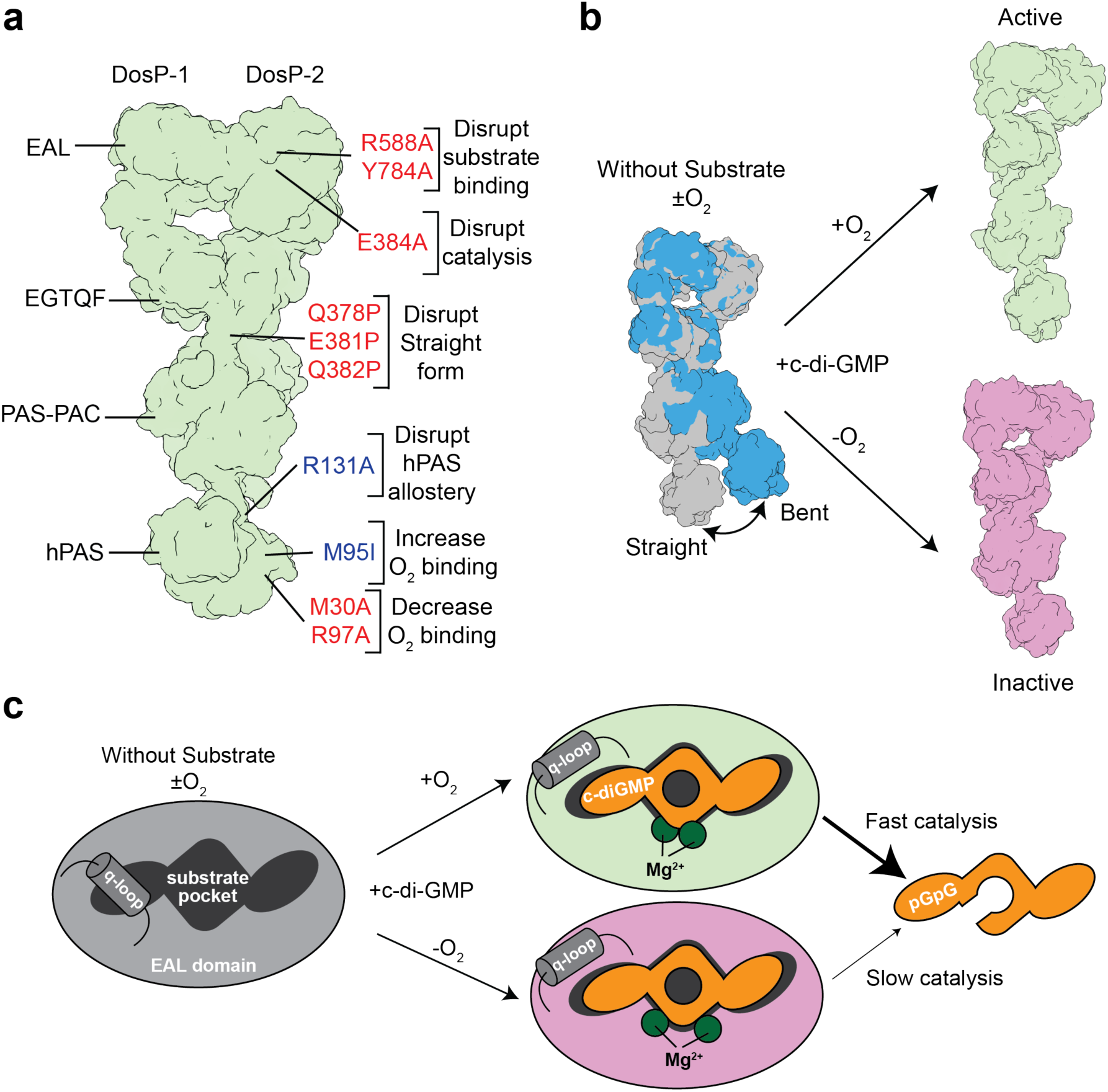
Model for DosP allostery. **a,** Schematic of DosP derived from the atomic model for DosP^FLAG-WT^ bound to c-di-GMP. The two DosP monomers and domains are labeled. Sites interrogated through mutagenesis are highlighted with brackets. Mutations colored red reduce the catalytic activity of DosP, and mutations colored blue increase the catalytic activity of DosP. **b,** Model for DosP allostery. In the absence of substrate, DosP exists in two conformations. Substrate binding stabilizes an oxy-state ON or deoxy-state OFF form. **c,** Schematic for the state of the EAL domain in the oxy and deoxy-states without and with substrate.

While the simplest model might posit that DosP is constitutively active whenever the inhibitory signal is not sent from the hPAS domain, that is clearly not the full story. The mutations DosP^Q378P^, DosP^E381P^, and DosP^Q382P^ in the hinge helix region between the PAC-PAS and EGTQF domains resulted in proteins that were less active than DosP^WT^ despite retaining a normal O_2_ binding response (**Fig. 3d-e**). The proline substitutions would be expected to disrupt the straight conformation that is stabilized by c-di-GMP, and the low activity of these mutants suggest that the straight form is active whereas the bent form is inactive. The observation of deoxy-state behavior in O_2_-ligated proteins supports a role for the intervening domains in allosteric communication required for both the ON and OFF states. We also find that it is possible to extinguish the phosphodiesterase activity by weakening the affinity for O_2_ and thereby destabilizing the ON-state. This can be accomplished by disrupting the hPAS-hPAS dimer interface (M30A) or by removing the stabilizing hydrogen bond donor (R97A). Our structures of the fully oxy/oxy DosP show that the key residues of the F-G loop are supported by contacts from the other hPAS monomer (**Fig. 6h-i**), and experiments with DosP^M30A^ revealed the importance of the hPAS dimeric interface in the ligation of O_2_ (**Fig. 7a-b**). In contrast, DosP^R97A^ removes a side chain that stabilizes O_2_ binding to the heme and enables M95 to compete more effectively against O_2_ for heme binding. Through separate mechanisms, both the M30A and R97A substitutions decrease O_2_ binding and thereby decrease the phosphodiesterase activity (**Fig. 7f**). In principle, regulatory partners could influence DosP at several different points along the protein.

With the caveats that our structural experiments relied on DosP^R97A^ as a surrogate for the deoxy, OFF-state DosP, and that our ∼3-4 Å resolution of the catalytic domain precludes confident modeling of the fine details, we conclude the following regarding the regulation of DosP by O_2_. First, DosP is dimeric under both aerobic and anaerobic conditions. This differentiates DosP from other O_2_ sensor-effector proteins such as DevS, an O_2_-regulated kinase from *Mycobacterium tuberculosis* that has been shown to change oligomeric state upon transition between its oxy and deoxy-states ^32^. Second, in the absence of substrate, DosP exists in an equilibrium of the straight and bent forms, regardless of its heme status. We propose that, from an equilibrium of the bent and straight forms, c-di-GMP binding stabilizes a single substrate-bound ON (DosP^WT^) or OFF (DosP^R97A^) form (**Fig. 8b**). Given the high affinity for c-di-GMP (**Extended Data Figs. 6d and 8a**), it is plausible that DosP is largely found in the straight form and samples the bent form after product release. Third, and perhaps most importantly, our data suggest that substrate binding is not the O_2_-regulated step. The position of the q-loop did not change in DosP^WT^ and DosP^R97A^ without substrate; however, the loop was readily displaced from the catalytic pocket by the high-affinity binding of c-di-GMP in the substrate-bound structures of DosP^FLAG-WT^ and Dos^FLAG-R97A^ (**Figs. 4c-e and 6e**). We have not detected meaningful differences in the affinity for c-di-GMP between aerobic and anaerobic DosP^WT^ (**Extended Data Fig. 8a**), and c-di-GMP is well-resolved in our reconstruction of DosP^FLAG-R97A^ when the protein was vitrified in the presence of c-d-GMP (**Fig. 6e**). Moreover, while DosP^WT^ operates at its maximum rate until low nanomolar concentrations of c-di-GMP (**Fig. 1g**), DosP^R97A^ remains inactive in the presence of 100 µM substrate (**Fig. 6a**). Instead of mediating the regulation by O_2_, the q-loop may help to stabilize the EAL domain when substrate is absent and/or prevent the association of other types of cyclic dinucleotides.

Association of the c-di-GMP substrate stabilized both oxy- and deoxy-DosP, but the final states differed for the two forms. Our structure of DosP^WT^ revealed changes in the hPAS dimer interface resulting from the full ligation of O_2_, and biochemical experiments with the structure- guided mutants DosP^M30A^ and DosP^R131A^ identified critical roles for the hPAS dimer in regulating DosP activity. Subtle changes in the hPAS pocket result in large changes throughout the DosP dimer that ultimately resolve to very small changes in the substrate binding pocket (**Fig. 6f-k and Extended Data Fig. 9c-d**). We propose that the DosP OFF-state represents a block of the catalysis imposed by a shift of the catalytic metals rather than a barrier to substrate binding (**Fig. 8c)**. Our structural analysis revealed multiple conformational changes initiating from the hPAS domain and suggests how they propagate through the protein.

## Methods

### Plasmids

The parental expression vector was *pMcYED31*, in which the *E*. *coli malE* gene was fused translationally to the 5’ start codon of dosC in the *dosCP* operon ^1^. All constructs were confirmed by DNA sequencing of the open reading frame before their use for protein expression.

*Plasmids encoding N-terminal MBP-TEV-DosP and variants.* The *dosC*-coding sequence was excised from *pMcYED31* by site-directed mutagenesis to generate *pMcDosP*, where the *E*. *coli malE* gene became instead translationally fused to the 5’ start codon of *dosP*. Subsequently the TEV protease recognition sequence “ENLYFQM” was inserted between the *malE*- and the *dosP*-coding sequences by site-directed mutagenesis such that the final plasmid *pMcTEVDosP* could be used to produce MBP-TEV-DosP. Variants were generated by site-directed mutagenesis PCR of *pMcTEVDosP*.

*Plasmids encoding C-terminal 3xFLAG-tagged DosP and variants.* The *E*. *coli malE* gene was excised from the *pMcDosP* by site-directed mutagenesis PCR and then a 3xFLAG-coding sequence (<u>DYKDDDDK</u>GS<u>DYKDDDDK</u>GS<u>DYKDDDDK</u>) was inserted at the 3’ of the *dosP*-coding sequence to generate the plasmid *pPtacDosP3xFLAG*. This was used to express DosP^FLAG^. The *pPtacDosP3xFLA*G served as the parental template to generate *dosP* variants by site-directed mutagenesis.

### Protein Purification

#### Purification of DosP from MBP-TEV-tagged proteins

For protein production, 2 L of *E. coli* TG1 cells harboring pMcTEVDosP plasmids encoding MBP-TEV-tagged DosP^WT^ or DosP variants were grown at 37 °C in Luria Bertani (LB) broth supplemented with 200 µg/mL ampicillin and 0.20% (w/v) glucose, starting with a 0.2% (v/v) inoculum from an overnight culture. Once the culture reached an OD_600_ of about 0.45, expression was induced by the addition of isopropyl-D-1-thiogalactopyranoside (IPTG) to 0.50 mM, and expression was continued overnight at room temperature. The cells were harvested by centrifugation at 3,000 x g for 15 min at 4 °C. The pellet was resuspended in 100 mL ice-cold buffer M (50 mM Tris-HCl, 150 mM NaCl, 5.0 mM MgCl_2_, 2.0 mM DTT, 5 % (v/v) glycerol, pH 8.0) containing two protease inhibitor cocktail tablets (Roche) and 1mM phenylmethylsulfonyl fluoride (PMSF, Thermo Fisher Scientific). The resuspended cells were disrupted by 24 cycles of 5 s sonication at an amplitude of 50% and 5 s cooling. The lysate was clarified by ultracentrifugation at 503k x g for 1 h at 4 °C and the resulting supernatant was filtered through a 0.22 µm PES membrane.

The clarified lysate was mixed with 10 mL of Amylose Resin (New England BioLabs) that had been equilibrated with Column Buffer [50 mM Tris-HCl, 200 mM NaCl, 1.0 mM EDTA, 2.0 mM DTT, 5.0 % (v/v) glycerol, pH 8.0], incubated for 1 h at 4 °C with gentle agitation, and subsequently loaded to a gravity column and allowed to flow through. The column was washed three times with Column Buffer and then eluted with 15 mL Column Buffer supplemented with 10 mM maltose. The eluted protein was concentrated using a centrifugal filter unit (Millipore) and subjected to size exclusion chromatography (SEC) over a Superose 6 10/300 GL Increase SEC column (Cytiva) equilibrated with Column Buffer. Eluted fractions having pure MBP-TEV-tagged proteins, as determined from their absorption spectra, were combined, and stored at -80 °C until further use.

The purified MBP-TEV-tagged proteins were concentrated to about 80 µM with a centrifugal filter unit (Millipore) before adding 500 units of TEV protease (New England BioLabs) and incubating for 3 h at 23 °C. The TEV-cut proteins were subjected to gel filtration again, as described above. The eluted fractions with pure DosP, free of any tag, were combined and stored at -80 °C. Absorption spectra for the proteins in Column Buffer were recorded at 300-700 nm with a Cary 50 UV-Visible Spectrometer (Varian).

For the cryo-EM studies, the protein was purified in the same way except that the final SEC buffer lacked glycerol. The eluted fraction with the highest concentration was selected for cryo-EM grid preparation.

#### Purification of DosP^FLAG-WT^ and DosP^FLAG-R97A^

The plasmids encoding DosP^FLAG-WT^ or *DosP^FLAG-R97A^*were transformed into chemically-competent BL21 (DE3) cells. Transformants were selected on LB agar-ampicillin (100 µg/ml) plates. A single colony was picked and grown overnight at 37 °C in LB broth supplemented with 100 µg/ml ampicillin. Protein overexpression was carried out in four liters of LB broth medium supplemented with glucose 0.2% (w/v) and ampicillin (200 µg/ml), that were inoculated with 1% (v/v) overnight-grown primary culture. The culture was allowed to grow at 37°C while shaking at 220 rpm until the OD_600_ reached 0.4 before induction with 1 mM Isopropyl-D-1- thiogalactopyranoside (IPTG). Proteins were produced while culturing for an additional 20 hours at 25°C. Cells were harvested by centrifugation at 4000 x g for 10 mins at 4 °C. The pellet was resuspended in lysis buffer containing 50mM Tris-HCl pH 7.5, 150mM NaCl, 5mM MgCl_2_, 0.5 mM tris(2-carboxyethyl) phosphine (TCEP), 5% (v/v) glycerol and 1mM PMSF. For the cell lysis, sonication was performed for 5 min at an amplitude of 50%, using a 5-s on and 5-s off cycle for the entire duration. The lysate was centrifuged at 100,000 x g for 1 hour at 4 °C. Following centrifugation, M2 anti-FLAG affinity resin (Sigma-Aldrich) was added to the supernatant and allowed to bind while gently rocking for 1 h at 4°C. The solution was subsequently loaded onto a gravity column and allowed to flow through. The column was washed with 5 column volumes of buffer containing 50mM Tris-HCl pH 7.5, 200mM NaCl, 1mM EDTA, 0.5mM TCEP and 5% (v/v) glycerol. Bound proteins were eluted in column buffer supplemented with 400 µg/ml of FLAG peptides. The eluted protein was concentrated using a centrifugal concentrator and then subjected to SEC over a Superose 6 10/300 GL Increase column equilibrated with 50mM Tris-HCl pH-7.5, 200mM NaCl, 1mM EDTA, 1mM DTT. The eluted fraction with the highest concentration was selected for cryo-EM grid preparation.

### Phosphodiesterase assay

Anaerobic DosP^WT^, DosP^FLAG-WT^, or their variants were prepared by treating the protein solution (approximately 500 µL of 7 µM protein) with 2% (w/v) glucose together with 50 units of catalase and glucose oxidase in an anaerobic chamber (Coy, Grass Lake, MI). The DosP was added to a final concentration of 10 nM to buffer (20 mM Tris, 2.5 mM MgCl_2_, 2.0 mM DTT, pH 8.0) containing 2.0 µM c-di-GMP. At indicated times, 10-µL aliquots were withdrawn from the reaction and quenched with an equal volume of 10 mM EDTA, pH 8.0. The samples were heated at 95 °C for 5 min and then either directly quantified for c-di-GMP or stored at -20 °C for subsequent measurement. The c-di-GMP was quantified with a Cyclic-di-GMP Assay kit (Lucerna, Inc., Brooklyn, NY) following the manufacturer’s protocol. This assay uses a c-di-GMP-binding riboswitch to stabilize the Spinach aptamer and enable DFHBI-1T fluorescence. Briefly, 20 µL of each sample were added to a flat-bottom 96-well plate, then 180 µL of the c-di-GMP assay buffer containing the c-di-GMP sensor and fluorophore molecule was added to each sample well. The samples were covered with foil, incubated dark for 45-60 min at 23 °C and then read on a fluorescence plate reader (BioTek) with excitation of 469 nm and emission of 501 nm. The concentration of c-di-GMP in the samples was calculated from a c-di-GMP standard curve obtained alongside each experiment.

### Isothermal Titration Calorimetry

DosP was purified for isothermal titration (ITC) as described above except that the final SEC buffer consisted of 50mM Tris-HCl pH-7.5, 200mM NaCl, 2 mM CaCl_2_, and 1mM TCEP. ITC experiments were carried out in a Malvern iTC200 calorimeter at 30°C. DosP was included in the cell at a concentration of 20 µM and c-di-GMP was loaded into the syringe at a concentration of 200 µM. A first injection volume of 0.5 μL was employed, followed by 20 injections of 1.9 μL each; all injection rates were 0.5 μL/s. The stir rate was 750 rpm, and the reference power was 5.0 μcal/s. The time between injections was 120 s. During the acquisition, the data were treated on-the-fly with “adaptive filtering” in addition to the moving-average data filter with a period of 5 s. The data were integrated, analyzed, and illustrated using NITPIC ^33^, SEDPHAT ^34^, and GUSSI ^35,36^, respectively.

### Cryo-Electron Microscopy

#### Grid Preparation

Quantifoil Au 300 mesh R 1.2/1.3 grids were plasma cleaned with air at 30 mA for 80 seconds using a Pelco easiglow machine. 4 µL of protein samples at about 1 mg/mL were spotted onto the grids at 4°C in 100% humidity and blotted for 4.5 s followed by freezing in liquid ethane cooled by liquid N_2_, using a Vitrobot Mark IV. Grids were stored in liquid N_2_ until imaging. For substrate-bound samples, the protein solution was supplemented with 5 mM MgCl_2_ after SEC and then further supplemented with 100 µM c-di-GMP while on ice immediately before freezing (≤10 seconds).

#### Data collection

6,324 movies for the DosP^FLAG^ samples with c-di-GMP were collected at PNCC using a Titan Krios microscope operating at 300 kV, with the post-column energy filter (Gatan) and a K3 direct detection camera (Gatan), using SerialEM ^37^. Movies were acquired at a pixel size of 0.5305 Å in super-resolution mode. The total accumulated dose was 50 e-/Å^2^over 50 frames. The defocus range was -0.8 to -2.0 µm. 9 images were collected for each stage movement using the beam-tilt imaging method.

7,192 movies for DosP^R97A^ were collected at PNCC using a Titan Krios microscope operating at 300 kV, with the post-column energy filter (Gatan) and a K3 direct detection camera (Gatan), using SerialEM ^37^. Movies were acquired at a pixel size of 0.5328 Å in super-resolution mode. The total accumulated dose was 50 e-/Å^2^over 50 frames. The defocus range was -0.8 to - 2.0 µm. 9 images were collected for each stage movement using the beam-tilt imaging method.

11,358 movies for DosP^WT^ without c-di-GMP were collected at PNCC using a Titan Krios microscope operating at 300 kV, with the post-column energy filter (Gatan) and a K3 direct detection camera (Gatan), using SerialEM ^37^. Movies were acquired at a pixel size of 0.4133 Å in super-resolution mode. The total accumulated dose was 50 e-/Å^2^ over 50 frames. The defocus range was -0.6 to -1.6 µm. 18 images were collected for each stage movement using the beam-tilt imaging method to collect two movies per hole.

9450 movies for DosP^FLAG-R97A^ were collected at UT Southwestern using a Titan Krios microscope operating at 300 kC, with a Selectris Energy Filter and Falcon 4i direct detection camera (ThermoFisher) using SerialEM ^37^. Movies were acquired at a pixel size of 0.738 Å. The total accumulated dose was 57 e-/A^2^. The defocus range was -0.8 to -2.9 µm.

#### Image processing

Cryo-EM data processing was carried out using cryosparc v4.2 (all maps except DosP^FLAG(R97A)^) or cryosparc v4.4 (DosP^FLAG-R97A^) ^38^.

##### DosP^FLAG-WT^ with c-di-GMP

Imported movies were motion-corrected using Patch Motion Correction and Fourier-cropped to a pixel size of 1.061 Å (**Extended Data Fig. 7a**. Following Patch CTF estimation, a curated set of 4681 micrographs were selected for subsequent processing. Following rounds of initial manual picking and template picking to generate quality templates from 2D classes, 9.8 million particles were picked using template picking. For crude classification steps, particles were extracted at a pixel size of 2.65 Å. Particles were winnowed by two rounds of 2D classification and then by Heterogenous 3D refinement against ab-initio reconstructions from a subset of particles that represented one good class and two junk classes (**Extended Data Fig. 7b**). From this, 1.56 million particles were re-extracted at a pixel size of 1.35 Å. These particles were subjected to Non-Uniform 3D Refinement and yielded an “initial map” with a resolution of 3.7 Å (**Extended Data Fig. 7c**). This map and particles were used as the starting point to generate two maps with better resolved features for different regions of the protein. Following another Heterogenous 3D Refinement to select ∼1.1 million particles with better features for the C-terminal lobe, a mask containing these domains was generated and used in focused 3D Classification (**Extended Data Fig. 7d**). Three classes with well-resolved density for this region and the linker helix connecting to the middle domains were selected. These 307,809 particles were subjected to Non-Uniform Refinement, Local Motion Correction, and a final Non-Uniform Refinement to obtain a 3.21 Å map with well-defined features for the bound substrate and C-terminal domains. The same C-terminal mask was used for a final Local Refinement to yield a map of 3.11 Å resolution.

To generate a quality map for the hPAS and PAS-PAC domains, a mask covering the poorly-resolved N-terminal hPAS domains was generated and used in focused 3D classification of the particles from the consensus map. This classification yielded 157,857 particles with clear density for this domain (**Extended Data Fig. 7e**). These particles were subjected to Non-Uniform Refinement, Local Motion Correction, and a final Non-Uniform Refinement to obtain a 3.61 Å map for the full protein, which revealed the N-terminal domains. A mask covering the hPAS and PAS-PAC domains was used to carry out a local refinement and yielded a map of 3.80 Å resolution that had stronger features for this region (**Extended Data Fig. 7e**).

Local resolution estimates for the Local Refinement maps are provided in **Extended Data Fig. 7f**. The half maps from the final refinements were filtered using DeepEMhancer ^39^. Gold-standard FSC curves, Three-dimensional FSC curves, and viewing distribution plots for the maps of DosP^FLAG-WT^ + c-di-GMP are provided in **Extended Data Fig. 7g**. The maps represent the same global conformation when aligned in the 3.69Å map (“consensus map”) that represented the complete DosP (**Extended Data Fig. 7h**) and a composite map was generated in chimera X (**Extended Data Fig. 7i**). This map was used for model building and figure illustrations.

##### DosP^R97A^

Imported movies were motion-corrected using Patch Motion Correction and Fourier-cropped to a pixel size of 1.066 Å. Following Patch CTF estimation, a curated set of 7049 micrographs were selected for subsequent processing (**Extended Data Fig. 4a**). Following rounds of initial manual picking and template picking to generate quality templates from 2D classes, 10.5 million particles were picked using template picking. For crude classification steps, particles were extracted at a pixel size of 3.31 Å. Particles were winnowed by two rounds of 2D classification and then by Heterogenous 3D Refinement against ab-initio reconstructions from a subset of particles that represented one junk class and two distinct conformations of DosP^R97A^ (**Extended Data Fig. 4b-c**). Particles for the two good classes were re-extracted at a pixel size of 1.33 Å and subjected to another round of Heterogenous 3D Refinement against four maps corresponding to two copies of each good map from the first Heterogenous 3D Refinement. 353,471 particles corresponding to the “straight conformation” were selected and subjected to local motion correction and Non-Uniform 3D Refinement to yield a map of 3.65 Å. 391,512 particles corresponding to the “bent conformation” were selected and subjected to local motion correction and Non-Uniform 3D Refinement to yield a map of 3.43 Å. Gold-standard FSC curves, Three-dimensional FSC curves, and viewing distribution plots for both maps are provided in **Extended Data Fig. 4d**. For model building and illustrations, the half maps from the final refinements were filtered using DeepEMhancer ^39^.

##### DosP^WT^

Two datasets from the same batch of grids were collected using the same microscope at PNCC. Imported movies were motion-corrected using Patch Motion Correction and Fourier-cropped to a pixel size of 0.8266 Å (**Extended Data Fig. 2a-b)**. Following Patch CTF estimation, a curated set of 5,642 micrographs were selected from the first dataset and 5,526 micrographs were selected from the second dataset. Each data set was processed individually for template-based particle picking, 2D classification, and initial heterogenous 3D refinement to select subsets of good particles. Particles were extracted at a pixel size of 1.61 Å. For the first dataset, 2.6 million particles were picked by template picking and winnowed to 658K after one round to 2D classification to remove junk particles. For the second dataset, 2.4 million particles were picked by template picking and winnowed to 866K after one round of 2D classification to remove junk particles. Iterative Heterogenous 3D refinement was used to further winnow the particles. A combined subset of 680,540 particles were subjected to heterogenous 3D refinement against four maps (2 maps generated from each dataset). Two good classes were obtained, yielding 191,953 particles for the “straight conformation” and 154,173 particles for the “bent conformation.” The final particles were subjected to Non-Uniform 3D Refinement to generate reconstructions of 3.91 and 3.97 Å, respectively. The final maps were filtered using DeepEMhancer ^39^ and the resulting maps are colored by local resolution (**Extended Data Fig. 2c**). Gold-standard FSC curves, Three-dimensional FSC curves, and viewing distribution plots for both maps are provided in **Extended Data Fig. 2d**.

##### DosP^FLAG-R97A^ with c-di-GMP

Imported movies in eer format were subdivided into 55 frames. Following Patch CTF estimation, a curated set of 9,228 micrographs were selected. Initial blob picks yielded templates that were used to pick 2.5 million particles (**Extended Data Fig. 8b-d**). Following 2D classification to remove junk particles, 780K particles were used to generate 6 ab initio models and then subjected to heterogeneous refinement against those maps. A single good class, with 186,761 particles was obtained that showed good features for the full protein. The particles were subjected to Reference-Based Motion Correction and yielded a map of 3.48Å global resolution. From this map, masks were generated covering the N-terminal portion or the C-terminal portions, which were subjected to local refinement to yield maps of 3.2 and 3.1 Å resolution, respectively (**Extended Data Fig. 8d**). The final maps were filtered using DeepEMhancer ^39^ and the resulting maps are colored by local resolution (**Extended Data Fig. 8d**). The two Local Refinement maps were aligned to the consensus map and combined to generate the composite map used for model building and illustrations (**Extended Data Fig. 8e**). Gold-standard FSC curves, Three-dimensional FSC curves, and viewing distribution plots for both maps are provided in **Extended Data Fig. 8f**.

#### Model building and refinement

PDBs 4HU4, 1S66, and the alphafold 2 prediction for DosP (accessed via Uniprot P76129) were used as starting models. Starting models were subjected to rounds of manual building in Coot^40^ and ISOLDE^41^ and real-space refinement in Phenix^42^. Refinement statistics are provided in **Table 1**. For model building and illustrations, the half maps from the final refinements were filtered using DeepEMhancer^39^. Map-Model FSC plots and examples of models built into cryo-EM density are shown in **Extended Data Fig.3**.

## Acknowledgements

We thank Marzia Miletto for expert assistance collecting cryo-EM data at PNCC. We thank Zhe Chen and Yang Li of the Structural Biology Lab core facility at UTSW for additional assistance with cryo-EM collections and advice on data analysis. A portion of this research was supported by NIH grant U24GM129547 and performed at the PNCC at OHSU and accessed through EMSL (grid.436923.9), a DOE Office of Science User Facility sponsored by the Office of Biological and Environmental Research. DLK is supported by NIH grants R00GM141261 and R01GM155152. We thank Margaret Phillips for critical comments on the manuscript.

## Author Contributions

WW carried out protein purification and biochemical experiments. PK carried out protein purification and structural studies. CB and S-CT carried out ITC studies. HRB assisted with assay development. MAGG conceived and supervised the research, carried out biochemical experiments, and wrote the paper. DLK conceived and supervised the research, carried out structural studies, and wrote the paper.

## Data availability

All data needed to evaluate the conclusions in the paper are presented in the paper and/or Supplementary Information. The structures of DosP^WT^ straight, DosP^WT^ bent, DosP^R97A^ straight, DosP^R97A^ bent, DosP^FLAG-WT^ + c-di-GMP, and DosP^FLAG-R97A^ + c-di-GMP have been deposited into the PDB under the accession codes 9CDR, 9CE0, 9CDR, 9CE0, 9CMF, and 9CLO, respectively. The related maps have been deposited into the Electron Microscopy Data Bank (EMDB). The accession codes are EMD- 45485 (DosP^WT^ straight), EMD- 45489 (DosP^WT^ bent), EMD-44524 (DosP^R97A^ straight), EMD- 44646 (DosP^R97A^ bent), EMD- 45645, EMD-45646, EMD-45665, and EMD-45746 (DosP^FLAG-WT^ + c-di-GMP), and EMD-45667, EMD-45669, EMD-45670, EMD-45727 (DosP^FLAG-R97A^ + c-di-GMP). Additional data related to this paper are available upon reasonable request from the authors.

## References Cited

1. Tuckerman, J.R. et al. An Oxygen-Sensing Diguanylate Cyclase and Phosphodiesterase Couple for c-di-GMP Control. Biochemistry 48, 9764–9774 (2009).

2. Schmidt Andrew, J., Ryjenkov Dmitri, A. & Gomelsky, M. The Ubiquitous Protein Domain EAL Is a Cyclic Diguanylate-Specific Phosphodiesterase: Enzymatically Active and Inactive EAL Domains. Journal of Bacteriology 187, 4774–4781 (2005).

3. Tanaka, A., Takahashi, H. & Shimizu, T. Critical Role of the Heme Axial Ligand, Met95, in Locking Catalysis of the Phosphodiesterase from Escherichia coli (Ec DOS) toward Cyclic diGMP*. Journal of Biological Chemistry 282, 21301-21307 (2007).

4. Shimizu, T. The Heme-Based Oxygen-Sensor Phosphodiesterase Ec DOS (DosP): Structure-Function Relationships. Biosensors 3, 211–237 (2013).

5. Delgado-Nixon, V.M., Gonzalez, G. & Gilles-Gonzalez, M.-A. Dos, a Heme-Binding PAS Protein from Escherichia coli, Is a Direct Oxygen Sensor. Biochemistry 39, 2685–2691 (2000).

6. Gilles-Gonzalez, M.A., Ditta, G.S. & Helinski, D.R. A haemoprotein with kinase activity encoded by the oxygen sensor of Rhizobium meliloti. Nature 350, 170–172 (1991).

7. Xing, J., Gumerov, V.M. & Zhulin, I.B. Origin and functional diversification of PAS domain, a ubiquitous intracellular sensor. Science Advances 9, eadi4517 (2023).

8. Yu, Z. et al. Gas and light: triggers of c-di-GMP-mediated regulation. FEMS Microbiology Reviews 47(2023).

9. Sousa, E.H.S. & Gilles-Gonzalez, M.-A. Chapter Five - Haem-Based Sensors of O2: Lessons and Perspectives. in Advances in Microbial Physiology, Vol. 71 (ed. Poole, R.K.) 235-257 (Academic Press, 2017).

10. Chang, A.L. et al. Phosphodiesterase A1, a Regulator of Cellulose Synthesis in Acetobacter xylinum, Is a Heme-Based Sensor. Biochemistry 40, 3420–3426 (2001).

11. Barends, T.R.M. et al. Structure and mechanism of a bacterial light-regulated cyclic nucleotide phosphodiesterase. Nature 459, 1015–1018 (2009).

12. Sousa, E.H.S., Tuckerman, J.R., Gonzalez, G. & Gilles-Gonzalez, M.-A. DosT and DevS are oxygen-switched kinases in Mycobacterium tuberculosis. Protein Science 16, 1708–1719 (2007).

13. Ross, P., Mayer, R. & Benziman, M. Cellulose biosynthesis and function in bacteria. Microbiological Reviews 55, 35–58 (1991).

14. Sommerfeldt, N. et al. Gene expression patterns and digerential input into curli fimbriae regulation of all GGDEF/EAL domain proteins in Escherichia coli. Microbiology 155, 1318–1331 (2009).

15. Méndez-Ortiz, M.M., Hyodo, M., Hayakawa, Y. & Membrillo-Hernández, J. Genome-wide Transcriptional Profile of *Escherichia coli* in Response to High Levels of the Second Messenger 3′,5′-Cyclic Diguanylic Acid *. Journal of Biological Chemistry 281, 8090-8099 (2006).

16. Tuckerman, J.R., Gonzalez, G. & Gilles-Gonzalez, M.-A. Cyclic di-GMP Activation of Polynucleotide Phosphorylase Signal-Dependent RNA Processing. Journal of Molecular Biology 407, 633–639 (2011).

17. Fekete, F.J., Marotta, N.J., Liu, X. & Weinert, E.E. An O2-sensing diguanylate cyclase broadly agects the aerobic transcriptome in the phytopathogen Pectobacterium carotovorum. Frontiers in Microbiology 14(2023).

18. Tarnawski, M., Barends, T.R.M., Hartmann, E. & Schlichting, I. Structures of the catalytic EAL domain of the Escherichia coli direct oxygen sensor. Acta Crystallographica Section D 69, 1045–1053 (2013).

19. Park, Suquet, C., Satterlee, J.D. & Kang, C. Insights into Signal Transduction Involving PAS Domain Oxygen-Sensing Heme Proteins from the X-ray Crystal Structure of Escherichia Coli Dos Heme Domain (Ec DosH). Biochemistry 43, 2738–2746 (2004).

20. Kurokawa, H. et al. A Redox-controlled Molecular Switch Revealed by the Crystal Structure of a Bacterial Heme PAS Sensor*. Journal of Biological Chemistry 279, 20186–20193 (2004).

21. Gonzalez, G. et al. Nature of the Displaceable Heme-Axial Residue in the EcDos Protein, a Heme-Based Sensor from Escherichia coli. Biochemistry 41, 8414–8421 (2002).

22. El-Mashtoly, S.F., Nakashima, S., Tanaka, A., Shimizu, T. & Kitagawa, T. Roles of Arg- 97 and Phe-113 in Regulation of Distal Ligand Binding to Heme in the Sensor Domain of Ec DOS Protein: RESONANCE RAMAN AND MUTATION STUDY*. Journal of Biological Chemistry 283, 19000–19010 (2008).

23. Dunham, C.M. et al. A Distal Arginine in Oxygen-Sensing Heme-PAS Domains Is Essential to Ligand Binding, Signal Transduction, and Structure. Biochemistry 42, 7701–7708 (2003).

24. Lechauve, C. et al. Heme Ligand Binding Properties and Intradimer Interactions in the Full-length Sensor Protein Dos from Escherichia coli and Its Isolated Heme Domain. Journal of Biological Chemistry 284, 36146–36159 (2009).

25. Navarro, M.V.A.S., De, N., Bae, N., Wang, Q. & Sondermann, H. Structural Analysis of the GGDEF-EAL Domain-Containing c-di-GMP Receptor FimX. Structure 17, 1104–1116 (2009).

26. Qi, Y. et al. Binding of Cyclic Diguanylate in the Non-catalytic EAL Domain of FimX Induces a Long-range Conformational Change *. Journal of Biological Chemistry 286, 2910–2917 (2011).

27. Yadav, M., Pal, K. & Sen, U. Structures of c-di-GMP/cGAMP degrading phosphodiesterase VcEAL: identification of a novel conformational switch and its implication. Biochemical Journal 476, 3333–3353 (2019).

28. Tchigvintsev, A. et al. Structural Insight into the Mechanism of c-di-GMP Hydrolysis by EAL Domain Phosphodiesterases. Journal of Molecular Biology 402, 524–538 (2010).

29. Huai, Q., Colicelli, J. & Ke, H. The Crystal Structure of AMP-Bound PDE4 Suggests a Mechanism for Phosphodiesterase Catalysis. Biochemistry 42, 13220–13226 (2003).

30. Bellini, D. et al. Dimerisation induced formation of the active site and the identification of three metal sites in EAL-phosphodiesterases. Scientific Reports 7, 42166 (2017).

31. Yoshimura, T., Sagami, I., Sasakura, Y. & Shimizu, T. Relationships between Heme Incorporation, Tetramer Formation, and Catalysis of a Heme-regulated Phosphodiesterase from Escherichia coli: A STUDY OF DELETION AND SITE- DIRECTED MUTANTS*. Journal of Biological Chemistry 278, 53105–53111 (2003).

32. Lobão, J.B.d.S., et al. Oxygen triggers signal transduction in the DevS (DosS) sensor of Mycobacterium tuberculosis by modulating the quaternary structure. The FEBS Journal 286, 479–494 (2019).

33. Keller, S. et al. High-Precision Isothermal Titration Calorimetry with Automated Peak-Shape Analysis. Analytical Chemistry 84, 5066–5073 (2012).

34. Houtman, J.C.D. et al. Studying multisite binary and ternary protein interactions by global analysis of isothermal titration calorimetry data in SEDPHAT: Application to adaptor protein complexes in cell signaling. Protein Science 16, 30–42 (2007).

35. Brautigam, C.A., Zhao, H., Vargas, C., Keller, S. & Schuck, P. Integration and global analysis of isothermal titration calorimetry data for studying macromolecular interactions. Nature Protocols 11, 882–894 (2016).

36. Brautigam, C.A. Chapter Five - Calculations and Publication-Quality Illustrations for Analytical Ultracentrifugation Data. in Methods in Enzymology, Vol. 562 (ed. Cole, J.L.) 109-133 (Academic Press, 2015).

37. Mastronarde, D.N. Automated electron microscope tomography using robust prediction of specimen movements. Journal of Structural Biology 152, 36–51 (2005).

38. Punjani, A., Rubinstein, J.L., Fleet, D.J. & Brubaker, M.A. cryoSPARC: algorithms for rapid unsupervised cryo-EM structure determination. Nature Methods 14, 290–296 (2017).

39. Sanchez-Garcia, R. et al. DeepEMhancer: a deep learning solution for cryo-EM volume post-processing. Communications Biology 4, 874 (2021).

40. Emsley, P., Lohkamp, B., Scott, W.G. & Cowtan, K. Features and development of Coot. Acta Crystallographica Section D 66, 486–501 (2010).

41. Croll, T. ISOLDE: a physically realistic environment for model building into low-resolution electron-density maps. Acta Crystallographica Section D 74, 519–530 (2018).

42. Afonine, P.V. et al. Real-space refinement in PHENIX for cryo-EM and crystallography. Acta Crystallographica Section D 74, 531–544 (2018).

