## Supplemental Data for "Structures of the multi-domain oxygen sensor DosP: remote control of a c-di-GMP phosphodiesterase by a regulatory PAS domain"

**Extended Methods and Data**

**Extended Data Methods:**

**Solution c-di-GMP binding assay**

Aerobic or anaerobic $DosP^{WT}$ (250 nM or 125 nM) equimolar with c-di-GMP substrate were equilibrated together for 15 min at 23°C in a final volume of 500 µL containing 20 mM Tris-HCl, 2.5 mM $CaCl_2$, 2 mM DTT, 500 nM ATP, pH 8.0. Note: ATP has no effect on DosP, and this nucleotide was included to prevent non-specific associations of the c-di-GMP with the spin-column membrane. The sample was loaded onto a spin column (Vivaspin 500, molecular weight cut-off of 100 kDa) that has been pre-equilibrated with the same buffer. An aliquot (20 µL) was removed from the column before spinning at 3000 x g for 15 s, which resulted in about 20 µL of flow through. The c-di-GMP concentration was determined using the c-di-GMP Assay Kit described under *Methods*. Total c-di-GMP was calculated from the mixture collected before the spin, and unbound c-di-GMP was calculated from the flow through after the spin.

**Extended Data Figures and Figure Legends:**

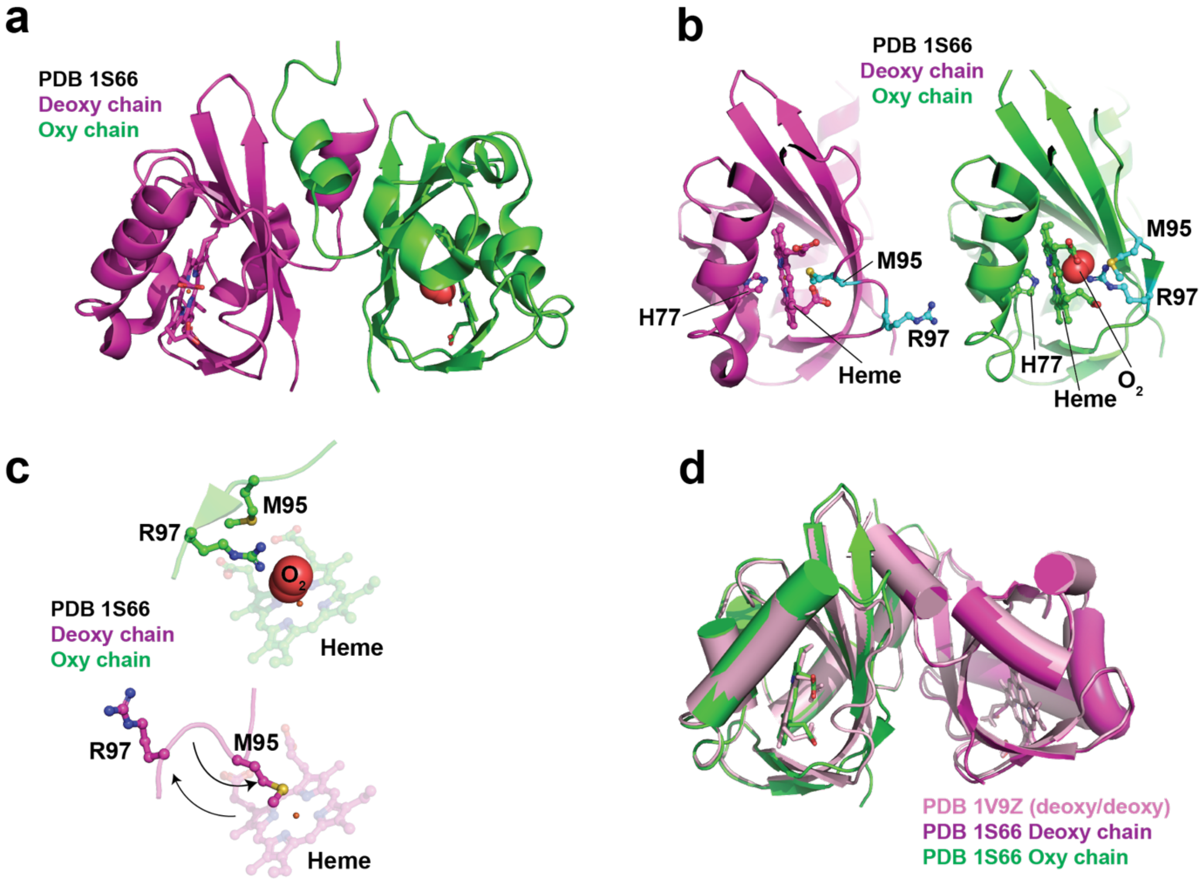


**Extended Data Figure 1.** Crystal structures of the oxy-state and deoxy-state DosP hPAS domain revealed the basis for internal competition between M95 and R97. **a)** PDB 1S66 is shown as a cartoon. The deoxy-state chain is colored magenta, the oxy-state chain is colored green. Heme is depicted as sticks and colored to match the cartoon. $O_2$ is depicted as red spheres. **b)** Side-by-side comparison of the separate chains of PDB 1S66. The chains were superposed and colored as in (a) except that the side chains of M95 and R97 are depicted as cyan sticks. H77 and heme are depicted as sticks and colored to match the cartoons. **c)** The heme groups and F-G loop residues including M95 and R97 from the two chains of PDB 1S66 are shown side-by-side from the same orientation. R97 and M95 are shown as sticks. $O_2$ is depicted as red spheres. Arrows indicate motion of R97 and M95 in the deoxy-state relative to the oxy-state. **d)** The mixed-state hPAS PBD 1S66 (magenta and green) is shown superposed with the deoxy-state hPAS PDB 1V9Z (pink).

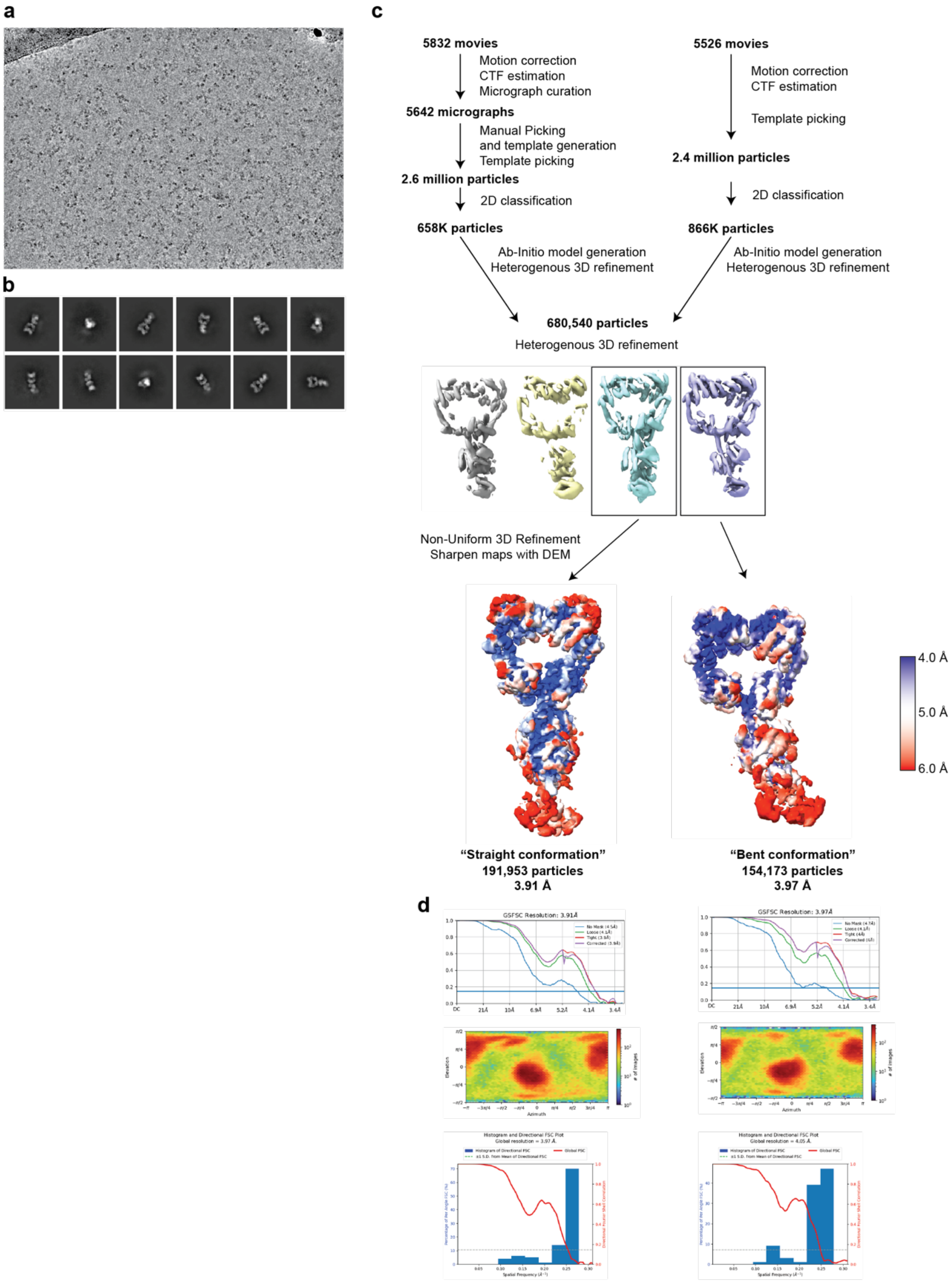


**Extended Data Figure 2.** Cryo-EM data processing for DosP$^{WT}$. **a)** Representative micrograph. **b)** 2D class averages. **c)** Data processing flow-chart. Final maps filtered using deepemhancer were colored by local resolution on a scale of 4Å to 6Å. **d)** FSC curves, viewing orientation maps, and directional FSC plots for the two maps.

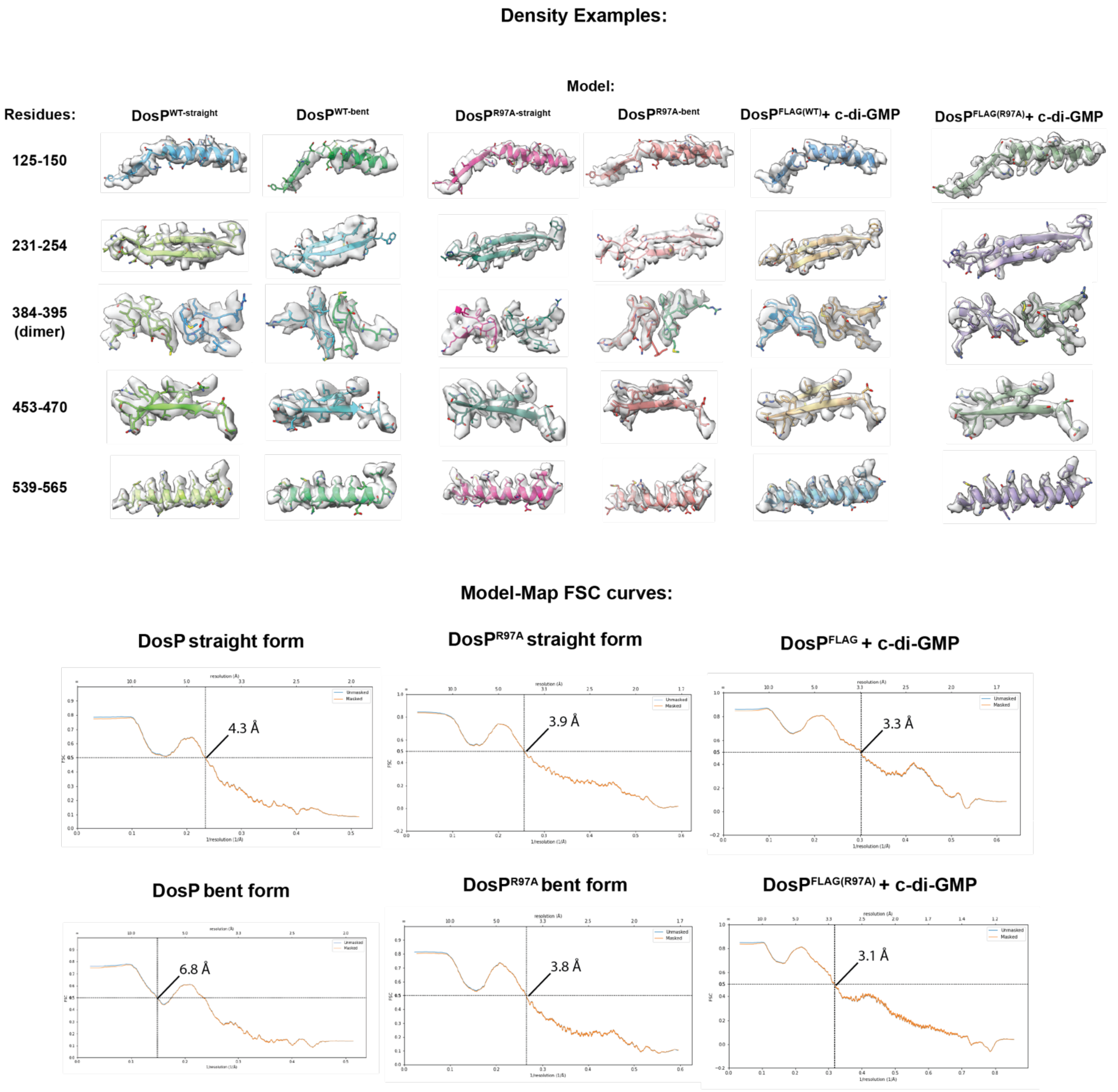


**Extended Data Figure 3.** Map-Model quality. (Top) Cryo-EM density maps and the atomic models in cartoon representation from the structures determined in this study. The sequence depicted in the figure is indicated on the left side of the map. (Bottom) Model-map FSC curves for all the structure solved in this study.

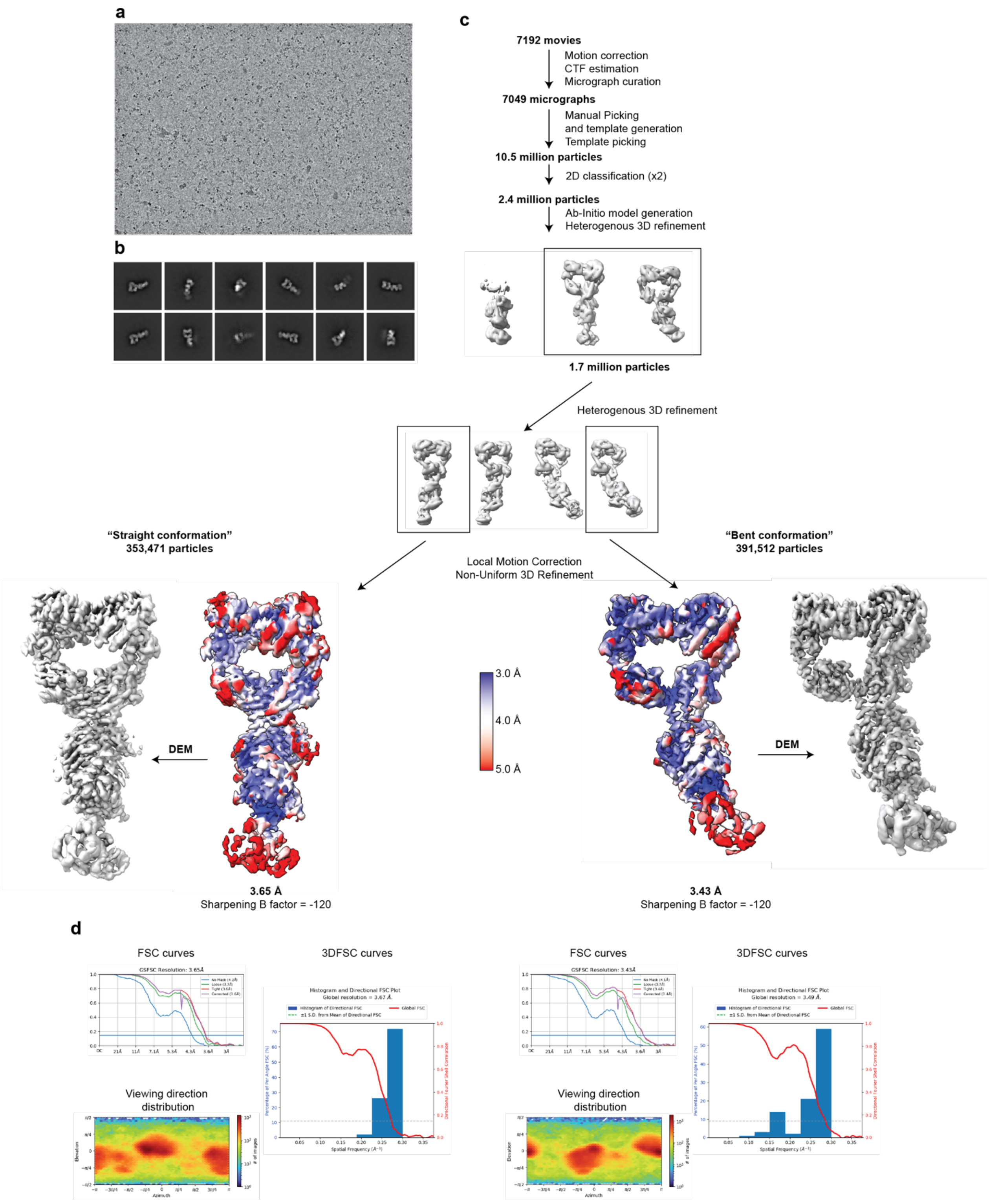


**Extended Data Figure 4.** Cryo-EM data processing for Dos$P^{R97A}$. **a)** Representative micrograph. **b)** 2D class averages. **c)** Data processing flow-chart. Maps sharpened with the indicated B-factor are colored by local resolution on a scale of 4Å to 5Å. Final maps filtered with deepemhancer are shown next to local resolution maps. **d)** FSC curves, viewing orientation maps, and directional FSC plots for the two maps.

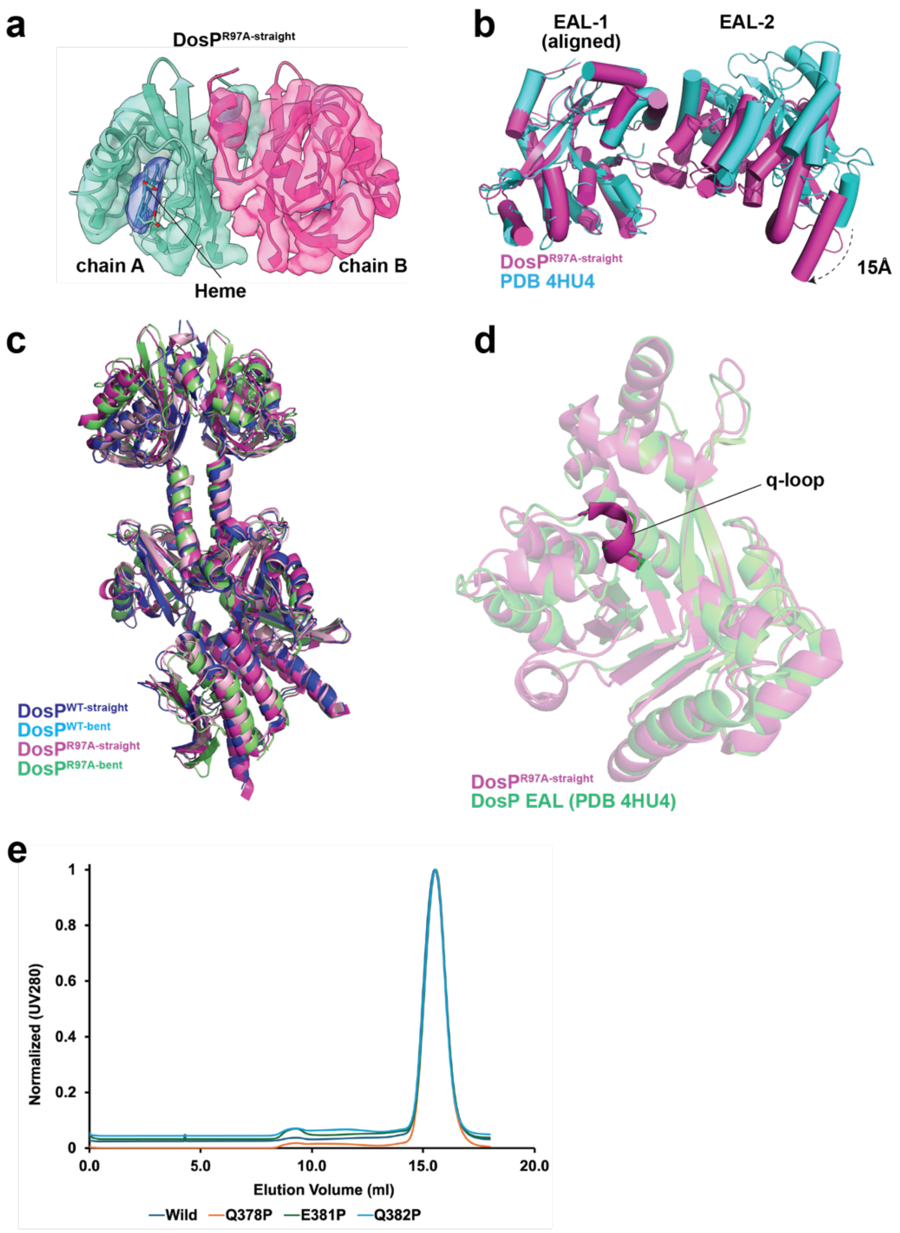


**Extended Data Figure 5. a)** The dimer structure of the h-PAS domain of DosP$^{R97A}$ in its straight form is depicted using both surface and cartoon representations, colored in green and magenta respectively, with a surface transparency set to 40%. The heme group is displayed in both surface and stick representations. The map density of the heme is colored blue, while the sticks are colored by heteroatoms. **b)** The dimeric form of the EAL domain of DosP$^{R97A}$ (magenta) is compared with the crystal structure of the DosP EAL domain (PDB ID: 4HU4), shown in cyan. The model is represented as cartoon cylindrical helices generated in PyMOL. The dimer structure was aligned to the EAL-1 domain. The black dashed line indicates the shift of the N-terminus of one monomer by approximately 15 Å. **c)** Cartoon representations show the superimposed structures of the bent and straight forms of DosP$^{WT}$ and DosP$^{R97A}$. DosP$^{WT}$ in its straight conformation is depicted in blue, while its bent conformation is shown in cyan. DosP$^{R97A}$ is represented in magenta, with its bent conformation displayed in green. **d)** The superimposed structure of DosP$^{R97A}$ straight form (magenta) with PDB 4HU4 shown in green. The q loop (residues 643-648) is depicted with 0% transparency, while the remainder of the cartoon representation is shown with 50% transparency. **e)** SEC traces for variants of DosP purified as described under *Methods*.

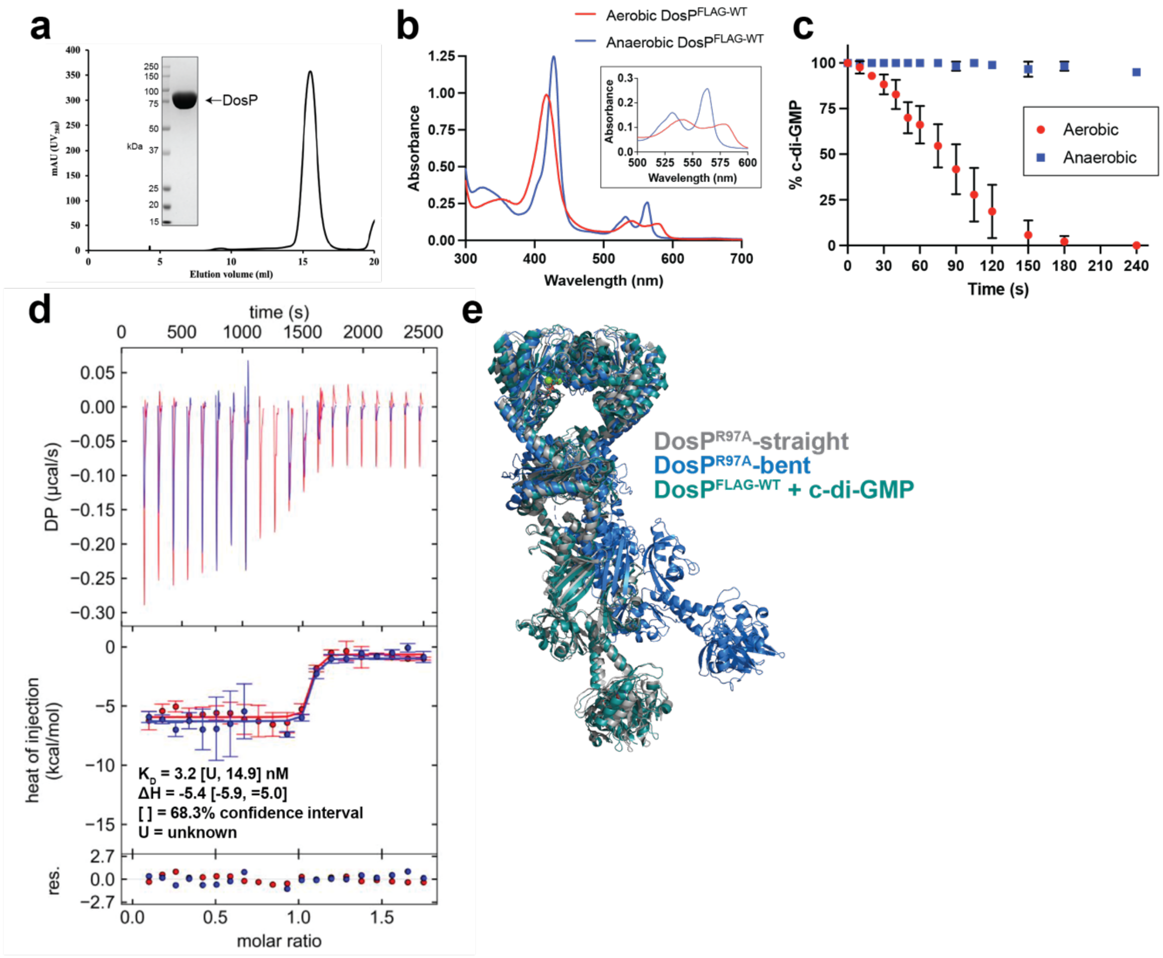


**Extended Data Fig. 6.** Characterization of $\text{DosP}^{\text{FLAG}}$ and substrate binding by DosP. **a)** Recovery of pure $\text{DosP}^{\text{FLAG-WT}}$ from a Superose 6 10/300 GL Increase SEC column after purification as described in *Methods*. Inset shows final sample collected from the major peak analyzed by Coomassie-stained SDS-PAGE. **b)** absorption spectra of $\text{DosP}^{\text{FLAG-WT}}$ under aerobic conditions (red), compared to anaerobic conditions (blue). **c)** Turnover of 2 μM c-di-GMP at 25°C by 10 nM aerobic $\text{DosP}^{\text{FLAG-WT}}$ (red circles), compared to anaerobic $\text{DosP}^{\text{FLAG-WT}}$ (blue squares), the assay buffer was 20 mM Tris-HCl, 2.5 mM $MgCl_2$, 2.0 mM DTT, pH 8.0. **d)** ITC analysis of DosP binding to c-di-GMP. The upper panel shows the SVD-corrected thermograms from NITPIC. The middle panel displays the integrated data points with their respective estimated error bars, and the lines results from the fit. The lower panel depicts the residuals between the data and the fit lines. **e)** Atomic models of $\text{DosP}^{\text{FLAG-WT}}$ (teal), $\text{DosP}^{\text{R97A}}$-straight (grey), and $\text{DosP}^{\text{R97A}}$-bent (blue) superposed through their EAL domains.

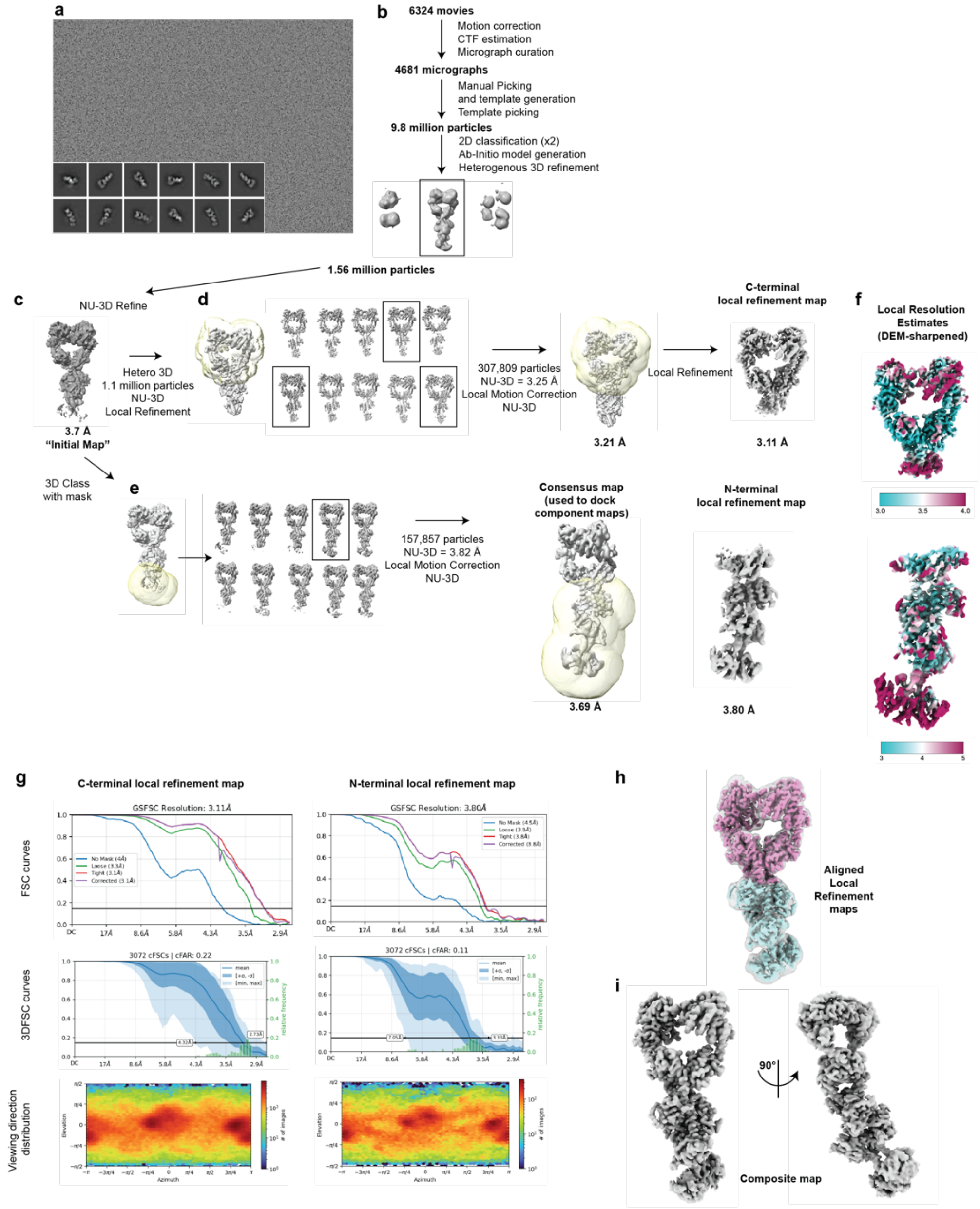


**Extended Data Figure 7.** Cryo-EM data processing for DosP$^{FLAG\text{-}WT}$ + c-di-GMP. **a)** Representative micrograph and 2D class averages. **b)** initial processing steps up to heterogenous 3D refinement based on *ab-initio* maps. **c)** The 3.7 Å global map used as the starting point for subsequent 3D classification without alignment. **d)** From left to right: masked 3D classification of particles based on their refined positions in (c). Boxed classes were selected for further refinement and local motion correction. The resulting 3.21A map and mask for final local refinement are shown next. Local-refinement of the C-terminal lobe yielded a 3.11 Å map. **e)** From left to right: masked 3D classification of particles based on their refined positions in (c). The boxed class was selected for further refinement and local motion correction. The resulting 3.69A map and mask of the N-terminal lobe for the final local refinement are shown next. Local-refinement yielded a 3.80 Å map with was filtered with deepemhancer and colored by local resolution from 3 to 6 Å. f) The final C-terminal (top) and N-terminal (bottom) maps were filtered using deepemhancer and colored by local resolution as indicated. **g)** FSC curves, viewing orientation maps, and conical FSC plots for the two maps. **h)** the two locally refined maps docked into the Consensus map. **i)** Composite map resulting from docking the N-term map and C-term map into the global map.

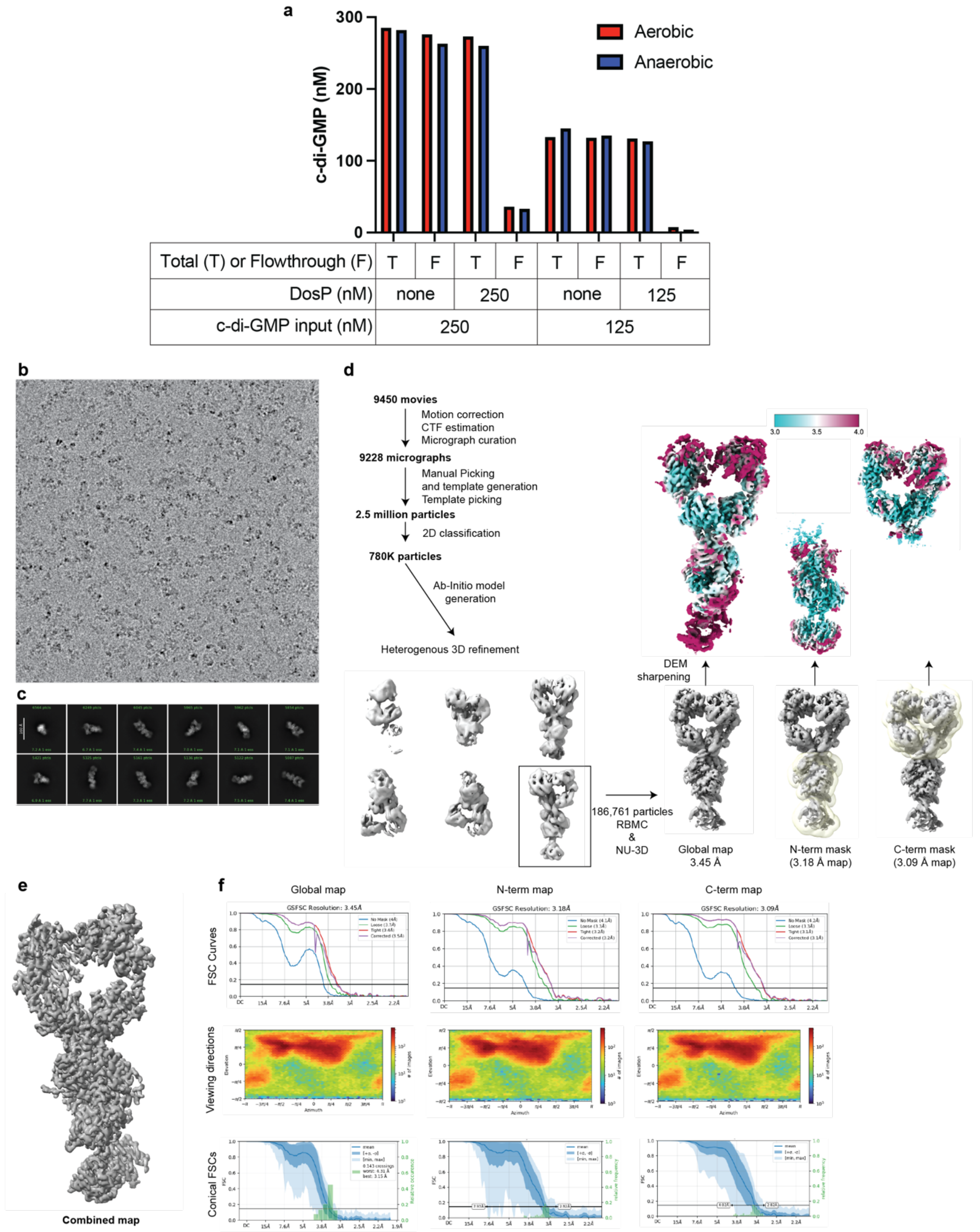


**Extended Data Figure 8.** Solution binding of c-di-GMP and cryo-EM data processing for DosP$^{FLAG-R97A}$ + c-di-GMP. **a)** Solution c-di-GMP binding assay. Levels of c-di-GMP were measured following equilibration at 25°C of 250 nM DosP$^{WT}$ in buffer ± 250 nM c-di-GMP, or equilibration of 125 nM DosP$^{WT}$ in buffer ± 125 nM c-di-GMP under aerobic (red) or anaerobic (blue) conditions. Following incubation, the samples were processed as described under *Extended Data Methods*. The assay buffer contained 20 mM Tris-HCl, 2.5 mM CaCl2, 2.0 mM DTT, 0.5 µM ATP, pH 8.0. **b)** Representative micrograph. **c)** 2D class averages. **d)** Data processing flow-chart. Final maps sharpened filtered with deepemhancer are shown colored by local resolution on a scale of 4Å to 5Å. **e)** Composite map resulting from docking the N-term map and C-term map into the global map. **f)** FSC curves, viewing orientation maps, and conical FSC plots for the two maps.

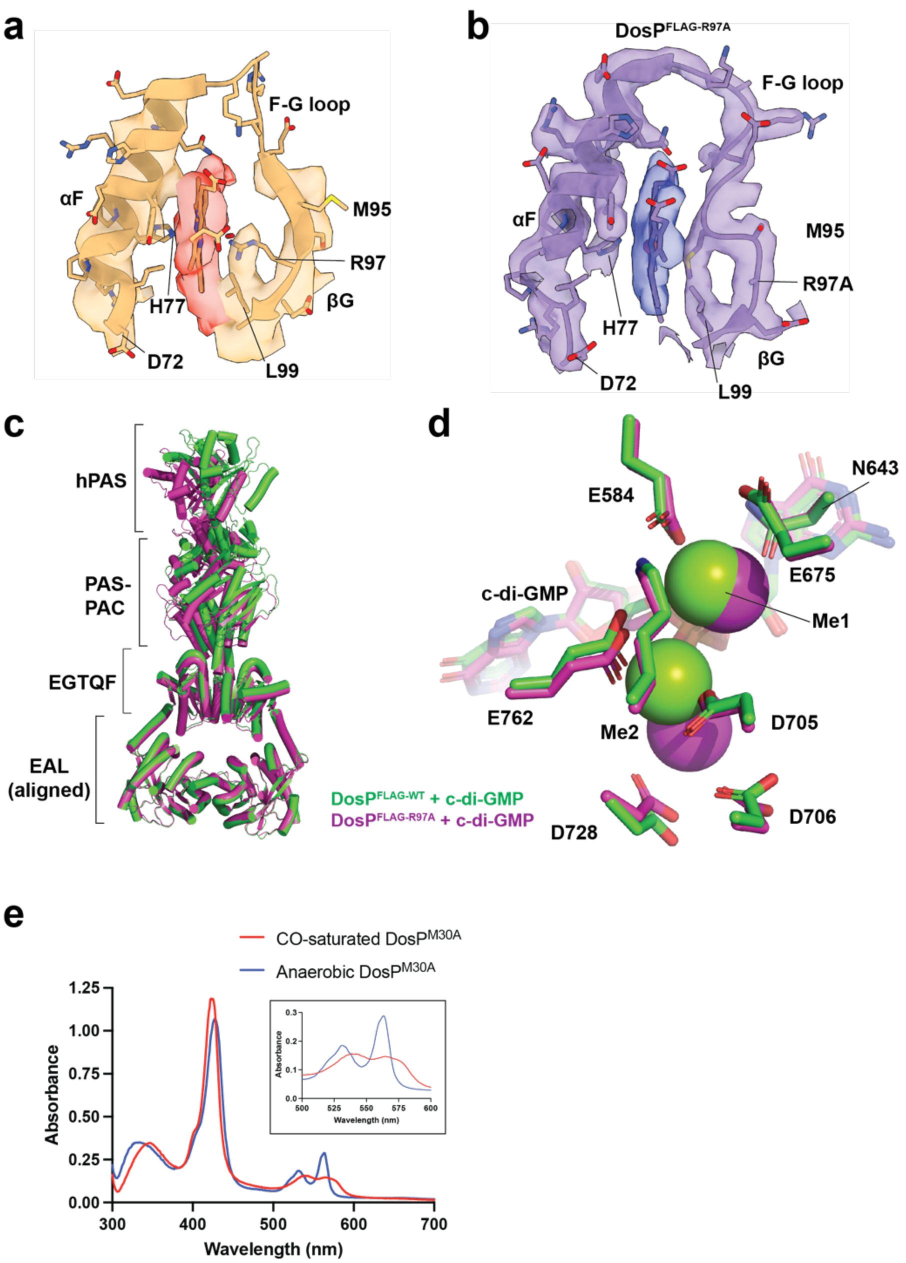


**Extended Data Figure 9: a)** DosP$^{FLAG-WT}$ (+ c-di-GMP) residues D72-L99 are shown in yellow cartoon with side chains and heme shown as sticks. Key residues are labeled. The map is shown at level 0.094 as transparent surface, carved at 3.0 Å. The map around the protein is transparent yellow and the map around the heme is transparent red. **b)** DosP $^{FLAG-R97A}$ (+ c-di-GMP) residues D72-L99 are shown in lavender cartoon with side chains and heme shown as sticks. Key residues are labeled. The map is shown at level 0.0959 as transparent surface, carved at 3.0 Å. The map around the protein is transparent lavender and the map around the heme is transparent blue. **c)** Atomic models for DosP$^{FLAG-WT}$ + c-di-GMP, shown in green, and DosP$^{FLAG-R97A}$ + c-di-GMP, shown in magenta, are superposed through the EAL domain. Domains are labeled with brackets. **d)** close-up of catalytic residues and magnesium atoms from one chain of DosP$^{FLAG-WT}$ and DosP$^{FLAG-R97A}$ colored as in (c). Side chains of metal-binding residues and c-di-GMP are shown as sticks. Magnesium metals shown as green or magenta spheres for DosP$^{FLAG-WT}$ and DosP$^{FLAG-R97A}$, respectively. **e)** Absorption spectra of DosP$^{M30A}$ saturated with CO (red), compared to anaerobic conditions (blue).
